# Infection-Induced Dynamin-2 Repurposing Drives Podosome Maturation and Mechanical Immunity

**DOI:** 10.64898/2026.07.01.735798

**Authors:** Chen Wang, Ting Chang, Tsung-Yu Lu, Ya-Wen Liu

**Affiliations:** Institute of Molecular Medicine, College of Medicine, National Taiwan University, Taipei, Taiwan

**Author notes:** Corresponding author: Ya-Wen Liu.

## Abstract

Macrophages are the vanguard of innate immunity, relying on phagocytosis to clear invading pathogens. Podosomes in macrophages have recently emerged as mechanical structures that assemble at phagocytic cups to facilitate pathogen clearance. However, how these unique actin structures are regulated to execute antimicrobial functions remains poorly understood. Here, we reveal that fungal infection drives podosome maturation to strengthen macrophage mechanical immunity. Exposure to the opportunistic fungus *Candida albicans* upregulates the scaffold protein Tks5 and activates Src kinase, which repurposes dynamin-2 from a membrane fission enzyme into an actin-bundling protein. This functional switch stabilizes the podosome actin core, enhancing the adhesion and mechanical force generation required to efficiently engulf, fold, and trap invading fungi. Ultimately, our findings define a mechanobiological paradigm in which infection-induced molecular repurposing drives macrophage antifungal immunity.

## Introduction

Immune cells internalize soluble ligands and small particles via endocytosis, whereas the engulfment of larger targets requires a mechanically distinct, actin-driven phagocytic process^1,2^. This challenge is most acute for elongated, rigid pathogens such as the hyphal form of *Candida albicans* (*C. albicans*), the main opportunistic and primary pathogenic fungus of humans^3,4^. To overcome these physical obstacles, immune cells must undergo dramatic cytoskeleton reorganization to generate the localized forces required to deform and confine invading pathogens.

Podosomes are dynamic, actin-rich protrusive structures that coordinate cellular adhesion and extracellular matrix (ECM) degradation, essential for cell migration^5–8^. In macrophages, they are abundant and well-recognized for extravasation and tissue infiltration^9–13^. Podosomes undergo rapid turnover, with their maturation into larger, stable structures driven by Src-activated Tks5, which recruits actin polymerization regulators and matrix metalloproteinases^14–16^, ultimately enhancing adhesion, matrix degradation, and protrusive force^17,18^. Beyond their classical role in motility, podosomes are increasingly recognized as mechanical structures during phagocytosis. Podosome-like structures have been observed at phagocytic cups and phagosomes, where they generate mechanical forces for target constriction, internalization, and phagosome maturation^19–28^. During *C. albicans* infection, podosomes have been implicated in the physical folding of hyphae within the phagosomes^26^. More recently, “phagocytic podosomes,” which share components and regulatory signals with classical ventral podosomes, were formally identified in primary human macrophages combating *Candida auris* (*C. auris*)^29^. Together, these studies support a mechanical role for podosomes during phagocytosis. However, the regulatory mechanisms driving this process, and whether podosome maturation is required to physically combat *C. albicans*, remain unknown.

Dynamin-2 (Dyn2) is a ubiquitous GTPase best known for assembling into membrane helices to drive GTP-dependent fission during endocytosis^30–32^. However, it also possesses the ability to crosslink filamentous actin (F-actin) into bundles within specialized actin structures, such as podosomes in muscle cells^33–35^ or stress fibers in podocytes^36^. This dual activity presents a biochemical paradox. *In vitro,* Dyn2-mediated actin bundling requires non-physiological low-salt conditions ([KCl]<75 mM), and GTP hydrolysis rapidly triggers the disassembly of Dyn-actin networks^33–35^. Consequently, it remains unclear how a GTPase optimized for rapid, seconds-long membrane remodeling during endocytosis^37,38^ can be converted into a stable structural component of podosomes, which persist for minutes^14,39,40^. We previously showed that the scaffold protein Tks5 directly recruits Dyn2 to podosomes^33^ and that Src-mediated phosphorylation of Dyn2 promotes podosome growth *in cellulo*^34^. Building on these findings, we hypothesize that the Tks5 interaction and Src phosphorylation coordinately act as a molecular switch to license the physiological actin-bundling activity of Dyn2.

Here, we identify an infection-responsive mechanism that promotes podosome maturation upon *C. albicans* infection. This process is driven by repurposing Dyn2 from a membrane fission enzyme to a cytoskeletal remodeler, thereby strengthening macrophage mechanical defense. These findings uncover a novel mechanobiological framework linking podosome maturation to macrophage mechanical immunity against fungal infection.

## Results

### *C. albicans* infection drives podosome maturation in BMDMs

Recent evidence suggests that podosomes function as mechanical structures during phagocytosis^24–29^. To determine whether pathogen infection modulates podosome formation, we monitored podosome morphology in primary mouse bone marrow-derived macrophages (mBMDMs) during *C. albicans* infection (Fig. EV1A). Confocal imaging showed that ventral podosomes present in resting macrophages (Fig. 1A i) rapidly disappeared in cells that had internalized yeast-form *C. albicans* at 0.5 hr post-infection (pi) (Fig. 1A ii, arrow), while neighboring cells lacking internalized fungi retained their podosomes (Fig. 1A ii, arrow head). This is consistent with previous reports demonstrating the relocation of podosome components to phagocytic cups or phagosomes during engulfment^23,28^. As infection progressed, intracellular fungal signals became increasingly diffuse, indicating progressive degradation, with most fungi degraded at 4 hr pi (Fig. 1A iii-iv). Strikingly, prominent podosome arrays assembled around the internalized fungi, encasing engulfed yeast (0.5 hr pi) and hyphae (2 hr pi) in structures highly colocalized with the podosome marker Cortactin (Fig. 1A v-viii).

**Fig. 1.**
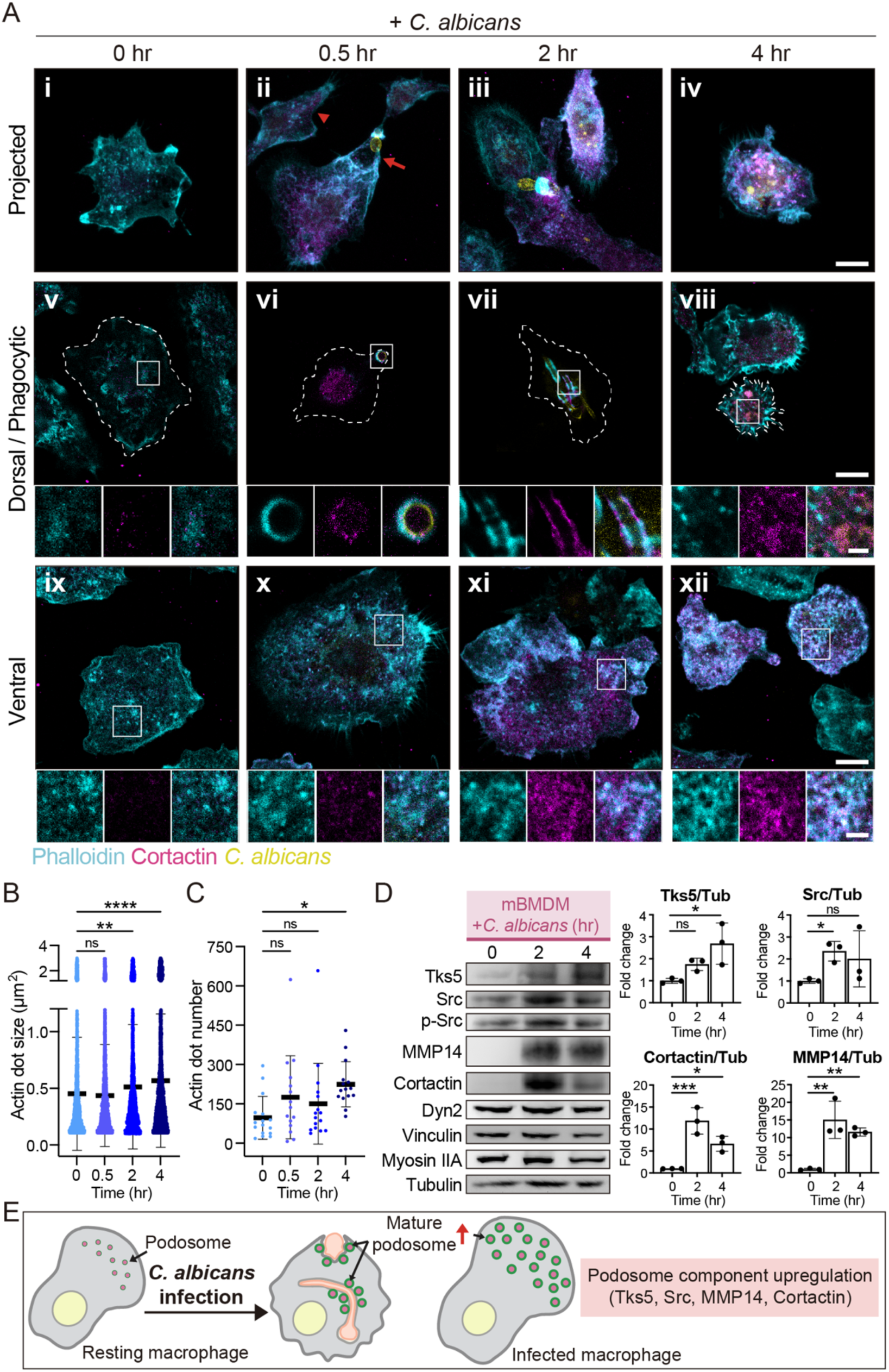
*C. albicans* infection promotes podosome maturation in mouse bone-marrow-derived macrophages. **(A)** Fluorescence imaging of mBMDMs infected with *C. albicans*. F-actin was labeled with phalloidin (cyan). Cortactin was stained with specific antibody (magenta). *C. albicans* was stained with CFW (yellow). Dashed lines outline the cell boundary. Scale bars, 10 μm and 2 μm. **(B, C)** Quantification of actin dot size (μm²) (n≥1551) and number per cell (n≥15) was analyzed with ImageJ. **(D)** Western blot and quantification of podosome components in mBMDMs infected with *C. albicans* for 0, 2, and 4 hours at MOI=5. Data are presented as mean ± SD of three independent experiments. Statistical significance was assessed using one-way ANOVA followed by Tukey multiple comparisons test (B, C) or the Fisher’s LSD test (D). ns, not significant; *, P<0.05; **, P<0.01; ***, P<0.001; ****, P<0.0001. **(E)** Schematic model of podosome maturation upon *C. albicans* infection.

To quantify infection-induced changes in podosome abundance and size, we analyzed macrophages exposed to *C. albicans* but lacking internalized fungi, using their retained ventral podosomes as a readout of global podosome dynamics. Infection led to a progressive increase in both podosome size and number per cell (Fig. 1A ix-xii, B, C), accompanied by elevated Cortactin levels and its enhanced accumulation at the podosome core (Fig. 1A ix-xii, insets), indicating widespread podosome maturation.

We next examined whether these morphological changes were accompanied by molecular signatures. Western blot analysis of infected mBMDMs revealed the significant upregulation of critical maturation drivers, specifically the scaffold Tks5, the matrix metalloproteinase MMP14 and Cortactin (Fig. 1D). The upstream tyrosine kinase Src was similarly elevated, with a trending increase in its active phosphorylated state (p-Src). Conversely, expression of other structural components, including Dyn2, Vinculin, and Myosin IIA, remained constant (Fig 1D). This molecular signature (upregulated Tks5/Src/MMP14) was conserved in RAW 264.7 macrophages (Fig. EV1B) and could also be triggered by bacteria or LPS treatment (Fig. EV1C, D). Together, these findings show that pathogen recognition induces a coordinated program of podosome maturation in macrophages (Fig. 1E).

### Tks5 promotes the actin-bundling activity of Dyn2 under physiological conditions

Dyn2 is an essential actin crosslinker required for podosome growth and maturation ^33–35^. Although its total expression level remained unchanged upon infection (Fig. 1D), immunofluorescence imaging revealed increased Dyn2 enrichment at ventral podosomes at 2 hr pi compared with 0.5 hr pi (Fig. 2A i, ii). Dyn2 also accumulated as patches and rings around actin structures surrounding engulfed hyphae (Fig. 2A iii), indicating enhanced recruitment to infection-induced podosomes. However, how Dyn2 transitions from a membrane-remodeling GTPase to a stable actin crosslinker within podosomes remains unclear.

**Fig. 2.**
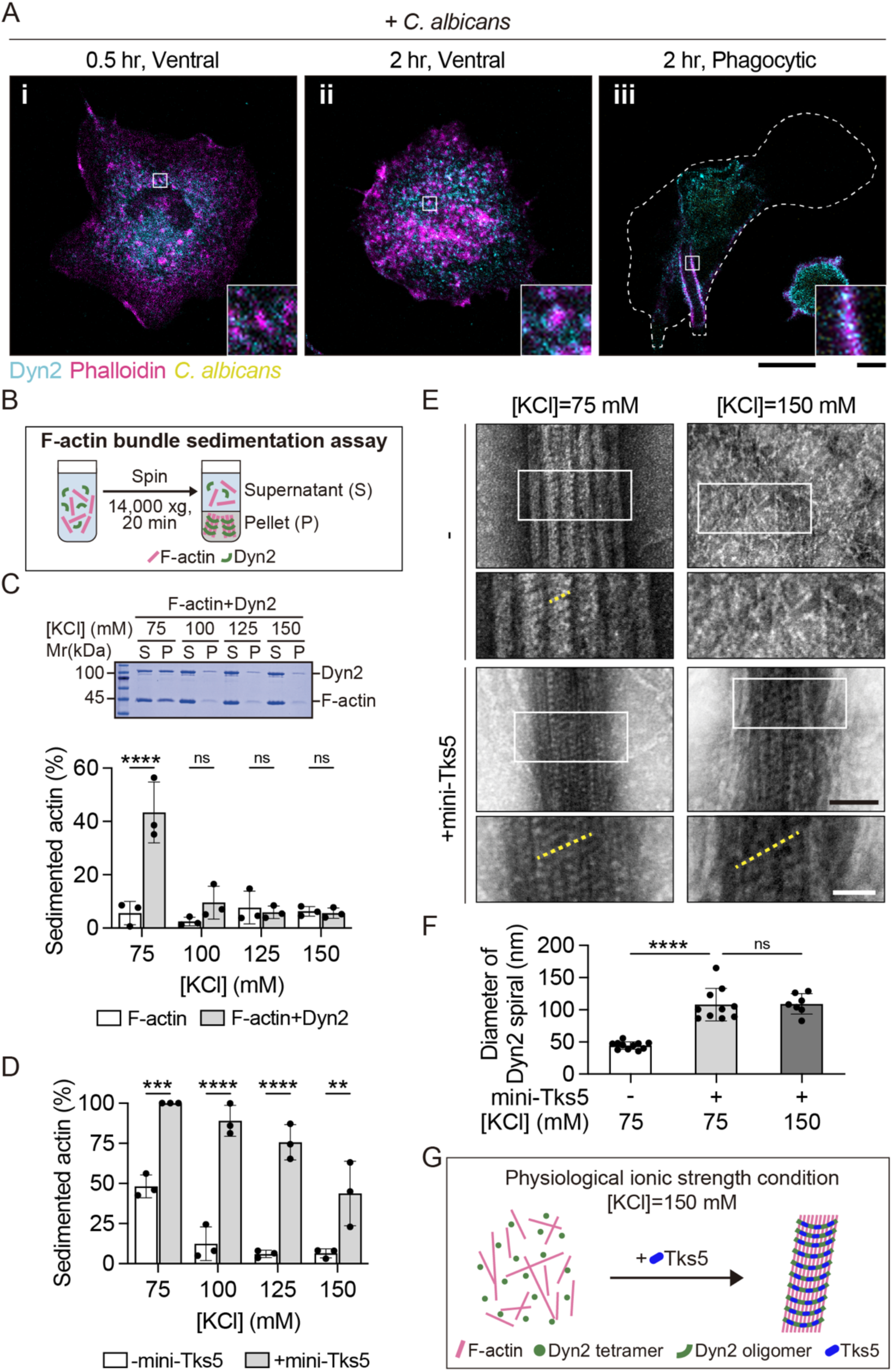
Tks5 facilitates Dyn2-mediated actin bundling under physiological conditions. **(A)** Fluorescence imaging of mBMDMs infected with *C. albicans*. F-actin was labeled with phalloidin (magenta). Dyn2 was stained with specific antibody (cyan). *C. albicans* was stained with CFW (yellow). Scale bars, 10 μm and 1 μm. **(B)** Schematic of the F-actin bundle sedimentation assay. **(C)** Representative SDS-PAGE gel and quantification of F-actin bundle sedimentation assay across a range of ionic strengths (75-150 mM KCl) in the presence or absence of Dyn2. **(D)** Quantification of actin bundling efficiency in the presence of mini-Tks5 across a range of ionic strengths (75-150 mM KCl). **(E)** Negative-stain TEM images showing Dyn2-actin bundling in the presence or absence of mini-Tks5, under buffer conditions containing either 75 or 150 mM KCl. Representative Dyn2 spirals are marked in yellow dashed line. Scale bars, black, 100 nm; white, 50 nm. **(F)** The diameters of Dyn2 spirals were measured using ImageJ (n≥7). Data are presented as mean ± SD from three independent experiments. Statistical significance was assessed using two-way (C, D) or one-way (F) ANOVA followed by Tukey multiple comparisons test. ns, not significant; **, P<0.01; ***, P<0.001; ****, P<0.0001. **(G)** Schematic model illustrating that Tks5 promotes Dyn2-actin bundling activity under near physiological ionic strength conditions.

Although dynamin assembles into spirals on membranes under physiological conditions^32^, membrane-free spiral formation occurs only at low ionic strength (<50 mM)^41^, indicating strong salt sensitivity of its oligomerization. We therefore tested whether its actin-bundling activity is similarly regulated using an F-actin sedimentation assay (Fig. 2B). Dyn2 efficiently bundled actin at 75 mM KCl but lost nearly all activity above 100 mM KCl (Fig. 2C). These findings indicate that Dyn2 actin-bundling activity is intrinsically incompatible with physiological ionic strength (∼150 mM), suggesting the existence of cellular mechanisms that enable its regulation *in vivo*.

We previously demonstrated that Tks5 directly binds and recruits Dyn2 to podosomes^33^; however, the mechanistic impact of Tks5 on Dyn2-mediated actin bundling under physiological ionic remains unclear. Given that the SH3A and SH3E domains of Tks5 directly interact with Dyn2 (Fig. EV2A, B), we evaluated the functional consequences of these interactions by introducing various recombinant Tks5 SH3 proteins into the F-actin sedimentation assay. While individual SH3A or SH3E domain failed to enhance bundling (and in some cases reduced it), the addition of a purified protein containing both domains (SH3A-E; hereafter referred to as ‘mini-Tks5’) or the complete SH3-domain scaffold (ΔPX) significantly increased actin sedimentation from ∼40% to ∼70% at 75 mM KCl (Fig. EV2C). Crucially, mini-Tks5 rescued Dyn2 bundling activity at physiological ionic strengths, where Dyn2 alone was inactive (Fig. 2D; Fig. EV2D). Negative-stain transmission electron microscopy (TEM) further showed that mini-Tks5 promoted the assembly of larger Dyn2 spirals around actin bundles at 75 mM KCl, increasing the spiral diameter from ∼50 nm to ∼100 nm (Fig. 2E, F). Remarkably, mini-Tks5 enabled comparable spiral formation at 150 mM KCl. Together, these findings identify Tks5 as an essential cofactor that licenses Dyn2 actin bundling under physiological salt conditions (Fig. 2G).

### Tks5 stabilizes Dyn2-actin bundles against GTP-induced disassembly

Because GTP hydrolysis drives Dyn2 disassembly from membranes^42,43^, we asked whether it similarly regulates Dyn2-actin bundles. Negative-stain TEM and F-actin sedimentation assays showed that GTP decreased the amount of sedimented actin (Fig. 3A; Fig. EV3A) and triggered Dyn2 dissociation from actin at 75 mM KCl (Fig. 3B). This effect was attenuated by mini-Tks5, which stabilized Dyn2-actin bundles, with partial protection also observed at 150 mM KCl (Fig. EV3B, C). To test the role of GTP hydrolysis in bundle disassembly, we utilized the non-hydrolyzable GTP analog GMPPCP. Dyn2 formed stable bundles at 75 mM KCl but not at 150 mM KCl in the absence of Tks5, whereas mini-Tks5 enabled stable assembly under both conditions (Fig. EV3D). These results indicate that Tks5 reinforces Dyn2-actin assemblies against GTP hydrolysis-mediated turnover.

**Fig. 3.**
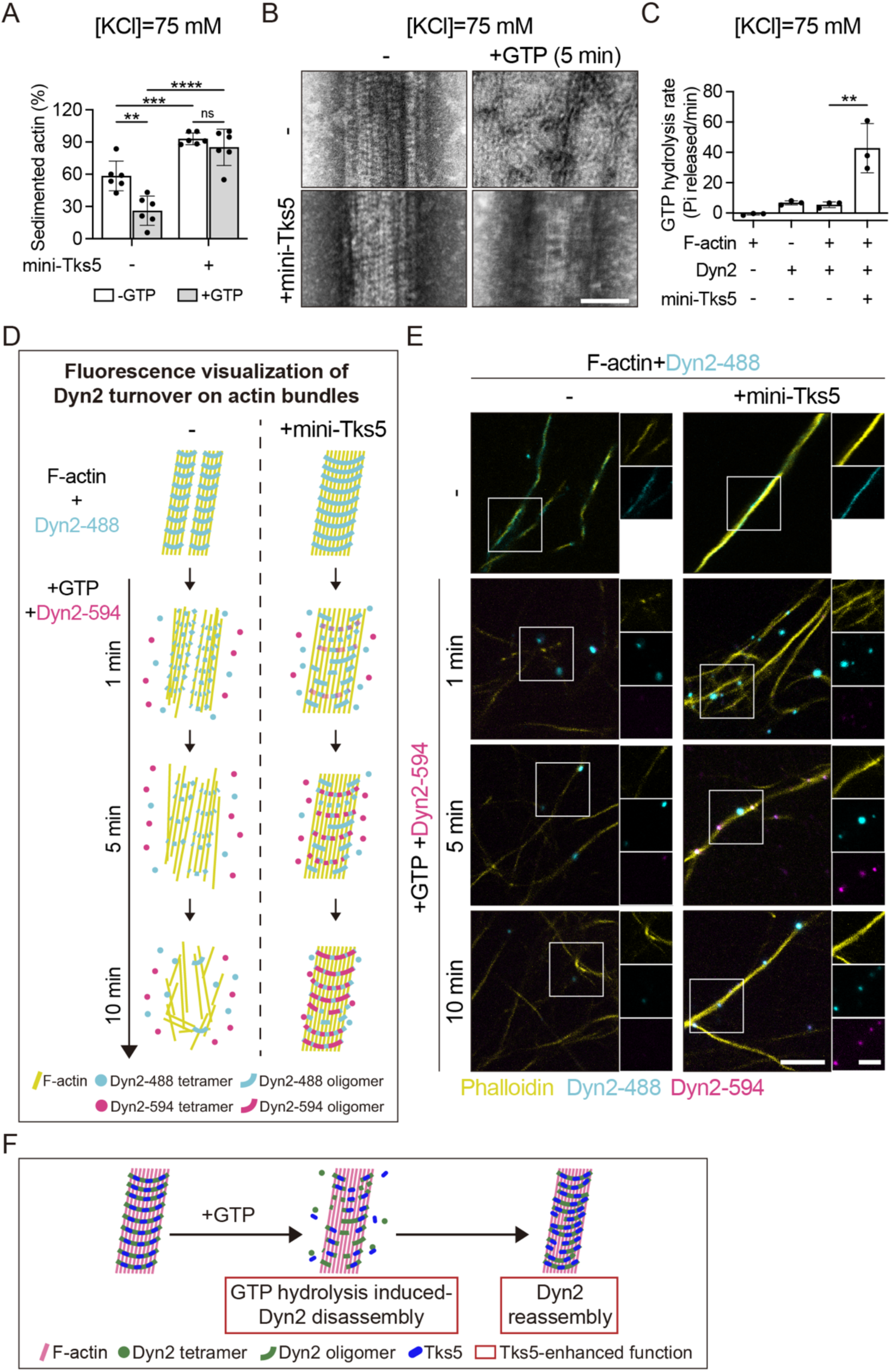
Tks5 sustains Dyn2-actin bundle stability against GTP-induced disassembly. **(A, B)** F-actin sedimentation assays (A) and negative-stain TEM images (B) of Dyn2-actin bundles incubated with or without mini-Tks5 and GTP for 5 min at 75 mM KCl. Bundling efficiency was quantified as the ratio of sedimented actin to total actin. Scale bar, 100 nm. **(C)** GTP hydrolysis rates of Dyn2-actin bundles in the presence or absence of mini-Tks5 at 75 mM KCl. Data are presented as mean ± SD from three independent experiments. Statistical significance was assessed using two-way (A) or one-way (C) ANOVA followed by Tukey multiple comparisons test. ns, not significant; **, P<0.01; ***, P<0.001; ****, P<0.0001. **(D, E)** Schematic workflow (D) and representative images (E) of visualizing Dyn2 turnover on actin bundles using two-color fluorescence-labeled Dyn2 (Dyn2-488 and Dyn2-594) in 75 mM KCl buffer. Scale bars, 5 µm and 2 µm. **(F)** Schematic model of Tks5-mediated reinforcement of Dyn2-actin bundles against GTP-induced disassembly.

We next examined whether Tks5 stabilizes bundles by inhibiting GTP hydrolysis activity of Dyn2. A malachite green assay showed that GTP hydrolysis rates of Dyn2 were unchanged in the presence of actin but were strongly increased by mini-Tks5, by ∼8-fold at 75 mM KCl (from 5.4 ± 2.1 to 42.8 ± 18.1 min^-1^) and ∼6-fold at 150 mM KCl (from 1.7 ± 0.3 to 10.1 ± 4.3 min^-1^) (Fig. 3C; Fig. EV3E). Thus, Tks5 does not inhibit Dyn2 turnover but instead promotes a highly active GTPase state. To further explore this dynamic behavior, we utilized two-color fluorescence-labeled Dyn2 to visualize molecular turnover directly on the bundles (Fig. 3D). Following GTP-induced bundle disassembly, mini-Tks5 promoted the rapid reassembly of both original Dyn2-488 (Dyn2 labeled with Alexa Fluor 488) and newly introduced Dyn2-594 (Dyn2 labeled with Alexa Fluor 594) onto the actin bundles, where they exhibited punctate enrichment (Fig. 3E).

Together, these findings show that Tks5 stabilizes Dyn2-actin assemblies not by suppressing GTP hydrolysis, but by promoting continuous turnover and rapid reassembly, thereby maintaining a dynamic yet mechanically stable actin network in podosomes (Fig. 3F). This kinetic “catch-and-release” cycle explains how a GTP-driven disassembly enzyme can be stably incorporated into the long-lived yet dynamic actin networks of mature podosomes.

### Dyn2-Y597 phosphorylation and Tks5 interaction cooperatively switch Dyn2 from membrane remodeling to actin bundling

Beyond its interaction with Tks5, we hypothesized that post-translational modification of Dyn2 regulates its actin-bundling activity. Dyn2 is phosphorylated by Src at Y597^44^, and a phospho-deficient mutant (Y597F) fails to localize to podosomes^34^, suggesting a role for this modification in Dyn2 function. However, how Y597 phosphorylation regulates Dyn2-actin bundling under physiological ionic conditions remains unclear. Using a phospho-mimetic mutant (Dyn2Y597E), we found that Y597 phosphorylation alone did not significantly alter actin bundling compared with Dyn2WT under either salt condition, with or without GTP (Fig. 4A; Fig. EV4A-C). However, in the presence of mini-Tks5, Dyn2Y597E exhibited markedly enhanced actin bundling, particularly under physiological ionic strength. These results indicate that Y597 phosphorylation and Tks5 interaction synergistically enhance Dyn2-actin bundling and stabilize bundles against GTP-induced dissociation under physiological ionic conditions.

**Fig. 4.**
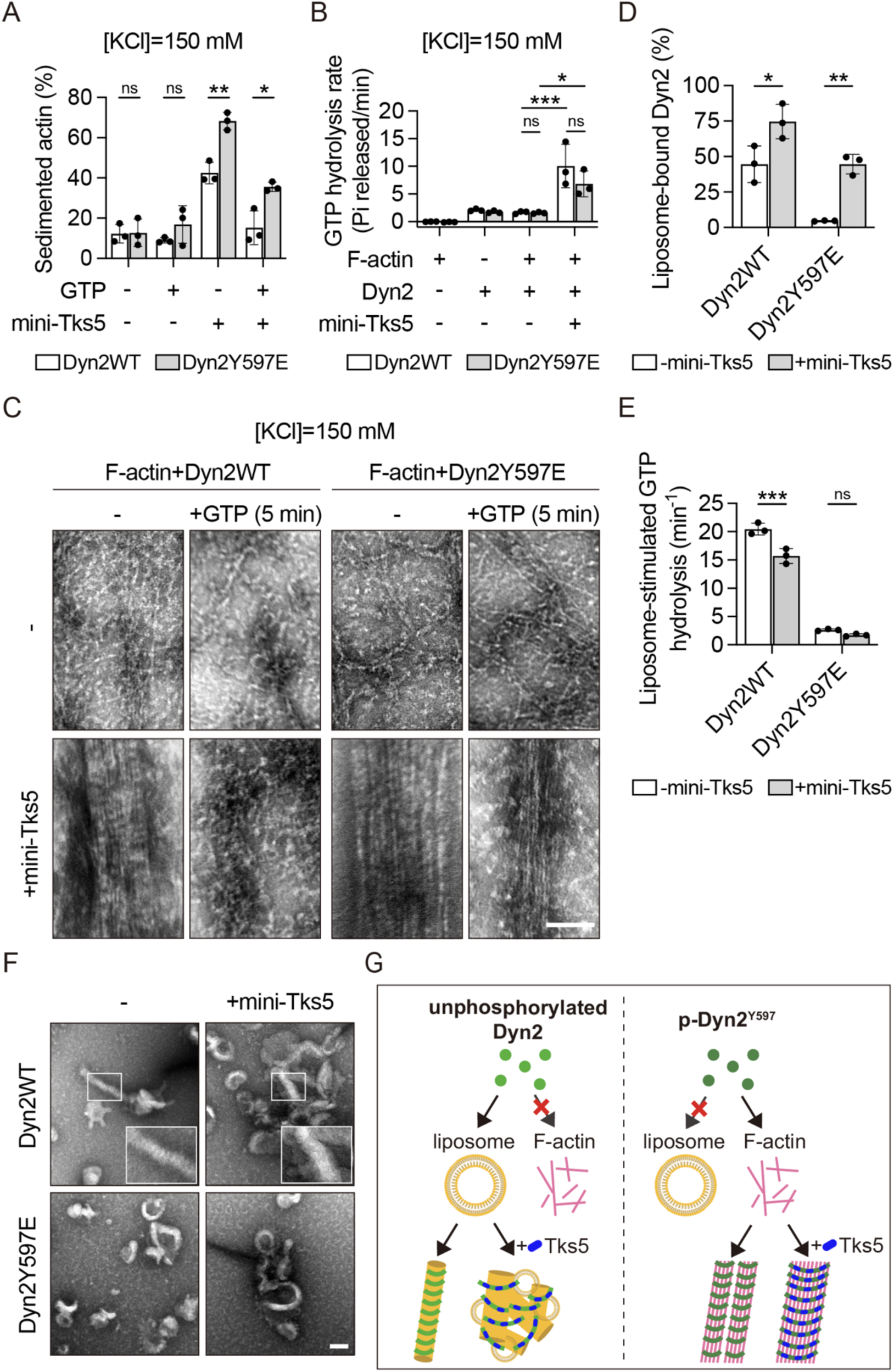
Dyn2-Y597 phosphorylation and Tks5 interaction synergistically promote Dyn2-actin bundling while diverting Dyn2 from membrane remodeling. **(A)** Quantification of F-actin sedimentation assays using Dyn2WT or Dyn2Y597E in 150 mM KCl buffer, with or without mini-Tks5 and GTP. **(B)** GTP hydrolysis rate of Dyn2WT- or Dyn2Y597E-actin bundles with or without mini-Tks5 in 150 mM KCl buffer. **(C)** Negative-stain TEM images of Dyn2WT- or Dyn2Y597E-mediated actin bundling with or without mini-Tks5 or GTP addition in 150 mM KCl buffer. Scale bar, 100 nm. **(D)** Liposome binding assay. Proteins were incubated with 100 nm liposomes and partitioned into supernatant (S, unbound) and pellet (P, bound) fractions via centrifugation. **(E)** Liposome-stimulated GTPase activity measured by malachite green assay. **(F)** Negative-stain TEM images of Dyn2 and/or mini-Tks5 incubating with liposomes. Scale bar, 100 nm. Data are presented as mean ± SD from three independent experiments. Statistical significance was assessed using two-way ANOVA followed by Tukey multiple comparisons test. ns, not significant; *, P<0.05; **, P<0.01; ***, P<0.001. **(G)** Schematic model illustrating that Tks5 and Y597 phosphorylation cooperatively drive Dyn2 toward actin bundling.

Similarly, mini-Tks5 increased GTP hydrolysis of Dyn2Y597E on actin bundles, as observed for Dyn2WT (Fig. 4B; Fig. EV4D). Negative-stain TEM showed that Dyn2Y597E formed stable actin bundles with mini-Tks5 under both salt conditions, even in the presence of GTP or GMPPCP (Fig. 4C; Fig. EV4E, F). Consistently, fluorescence imaging revealed that while GTP induced dissociation of Dyn2WT and Dyn2Y597E from actin, mini-Tks5 prevented this dissociation and further enhanced Dyn2 recruitment (Fig. EV5A, B). Notably, Dyn2Y597E showed stronger actin association than Dyn2WT, identifying Y597 phosphorylation as a priming event that stabilizes Dyn2-actin assemblies.

We further examined how mini-Tks5 influences Dyn2-lipid interactions. Interestingly, mini-Tks5 enhanced the liposome-binding ability of both Dyn2WT and Dyn2Y597E (Fig. 4D; Fig. EV5C). However, both the presence of mini-Tks5 and Y597 phosphorylation severely inhibited the liposome-stimulated GTPase activity of Dyn2 (Fig. 4E). Negative-stain TEM imaging revealed that while Dyn2WT alone assembled into canonical spiral structures and tubulated liposome, the addition of mini-Tks5 caused dynamin-liposome clustering without spiral formation (Fig. 4F). For Dyn2Y597E, only liposome-like vesicular structures were observed. These findings suggest that Y597 phosphorylation acts as a molecular switch, directing Dyn2 to preferentially bind and crosslink actin rather than remodel the plasma membrane (Fig. 4G).

To further characterize the architecture of these assemblies, we performed cryo-electron microscopy (cryo-EM) on Dyn2-actin bundles formed in the presence of mini-Tks5 under 75 mM KCl (Fig. EV6A). Although sample heterogeneity precluded a high-resolution 3D reconstruction, 2D class averages showed Dyn2 organized into helical structures encircling the actin filaments (Fig. EV6B). These observations are consistent with our negative-stain TEM data and support a structural model in which Dyn2 assemble externally around the actin bundles.

### Dyn2-Y597 phosphorylation and Tks5 interaction strengthen Dyn2-actin bundles

Because Dyn2-actin bundles appeared thinker and straighter in the presence of mini-Tks5 (Fig. 3E; Fig. EV5A), we investigated whether Tks5 interaction and Y597 phosphorylation alter the mechanical resilience of these bundles. To assess bundle stiffness, we measured the persistence of the Dyn2-actin bundles by calculating the ratio of the end-to-end distance to the total contour length, adapting a metric commonly used to assess directional persistence of cell migration^45^. At 75 mM ionic strength, mini-Tks5 increased bundle persistence for both Dyn2WT and Dyn2Y597E (Fig. 5A, B). At 150 mM, bundles were barely detectable unless mini-Tks5 was added, consistent with our earlier findings. Notably, at this physiological ionic strength, Dyn2Y597E-actin bundles exhibited significantly greater persistence compared to Dyn2WT (Fig. 5C, D).

**Fig. 5.**
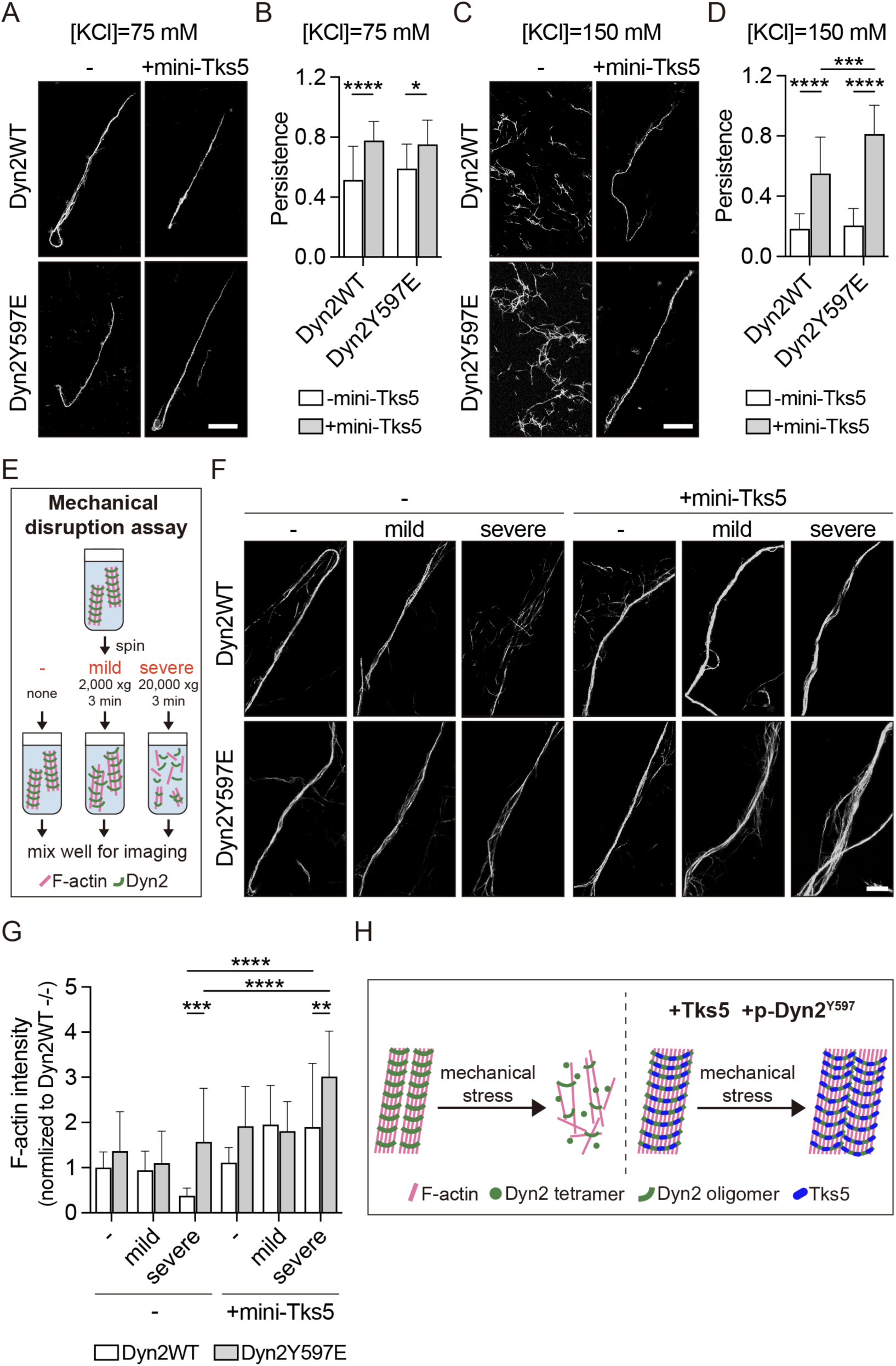
Dyn2-Y597 phosphorylation and Tks5 enhance the mechanical stability of Dyn2-actin bundles. **(A-D)** Fluorescence images of actin bundles formed by Dyn2WT or Dyn2Y597E, with or without mini-Tks5, stained with phalloidin in 75 and 150 mM KCl buffer. Bundle straightness was quantified by calculating the ratio of end-to-end distance to total contour length (n≥15). Scale bars, 10 µm. **(E)** Workflow of the mechanical disruption assay to assess the resistance of Dyn2-actin bundles to mechanical stress. **(F, G)** Representative fluorescence images of actin bundles assembled with Dyn2WT or Dyn2Y597E, with or without mini-Tks5, stained with phalloidin after mechanical disruption. Scale bar, 10 µm. Mechanical stability of the bundles assembled with Dyn2 was quantified by measuring mean phalloidin fluorescence intensity using ImageJ (n≥12). Data are presented as mean ± SD from three independent experiments. Statistical significance was assessed using two-way ANOVA followed by Tukey multiple comparisons test. *, P<0.05; **, P<0.01; ***, P<0.001; ****, P<0.0001. **(H)** Schematic model illustrating that Tks5 and Dyn2-Y597 phosphorylation synergistically reinforce the mechanical stability of actin bundles.

Furthermore, we tested whether Tks5 interaction and Dyn2-Y597 phosphorylation protect Dyn2-actin bundles from mechanical stress. Pre-formed Dyn2-actin bundles were subjected to varying degrees of shear force using different centrifugation speeds (Fig. 5E). While mild mechanical stress had little impact on Dyn2WT bundles, severe stress caused fragmentation and a marked reduction in F-actin signal. The presence of mini-Tks5, however, markedly preserved bundle integrity, as reflected by sustained F-actin signal and intensity (Fig. 5F, G). Strikingly, Dyn2Y597E-actin bundles displayed even higher intrinsic resilience, maintaining their structure even under severe disruption, a phenotype that was further stimulated by the addition of mini-Tks5. Together, these findings demonstrate that Tks5 interaction and Y597 phosphorylation synergistically reinforce Dyn2-actin bundles. This dual regulation not only increases bundling activity and mechanical strength but also enables the formation of larger, more cohesive bundle clusters under stress (Fig. 5H). Ultimately, this regulation may provide the structural resilience necessary for podosomes to maintain their integrity under immense mechanical challenges.

### Dyn2-Y597 phosphorylation and Tks5 expression promote macrophage podosome maturation

Building on our biochemical findings, we hypothesized that the structural strengthening of Dyn2-actin bundles by Y597 phosphorylation and Tks5 drives podosome maturation *in vivo*. To test this, we first generated RAW 264.7 macrophages stably overexpressing Tks5 (Tks5-OE, exhibiting a 7.1-fold increase) (Fig. 6A) and evaluated its podosome morphology. Tks5-OE macrophages exhibited a significant increase in podosome size while maintaining a total podosome number comparable to control cells (Fig. 6B-D). This indicates that the upregulation of Tks5, a phenomenon we previously demonstrated occurs naturally during *C. albicans* infection, is sufficient to drive macrophage podosome maturation.

**Fig. 6.**
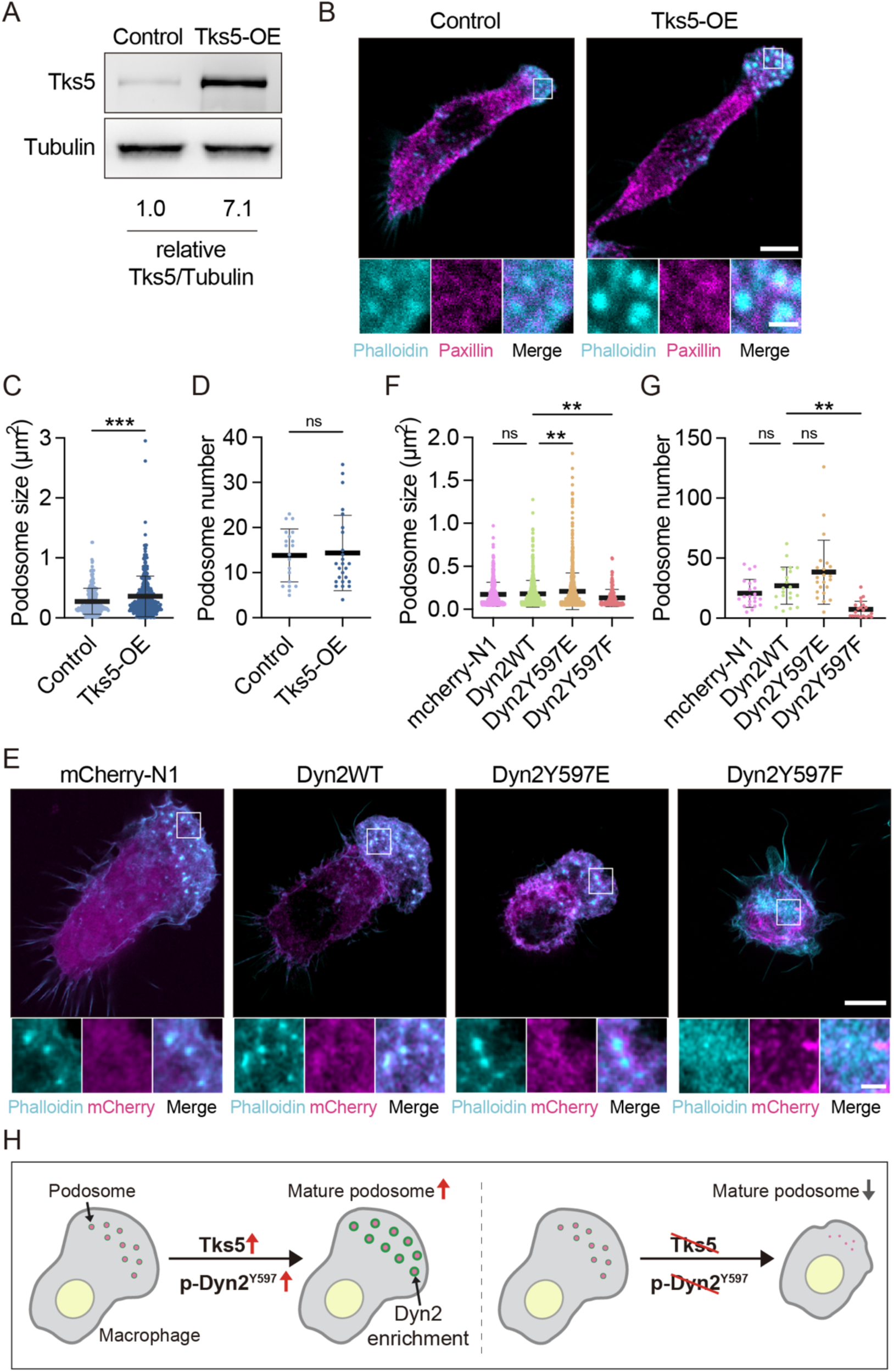
Dyn2-Y597 phosphorylation and Tks5 facilitate podosome maturation in macrophages. **(A)** Western blot analysis confirmed the overexpression of Tks5 in Tks5-OE cells compared to control cells. **(B)** Immunofluorescence staining of control and Tks5-OE RAW 264.7 cells. F-actin was labeled with phalloidin (cyan), and Paxillin was detected using specific antibodies (magenta). Scale bars, 5 μm and 1 μm. **(C, D)** Quantification of podosome size (μm²) (n≥277) and podosome number per cell (n≥20) were analyzed with ImageJ. **(E)** Fluorescence imaging of mCherry-N1 and DynWT-, Dyn2Y597E-, Dyn2Y597F-mCherry (magenta) transfected RAW 264.7 cells. F-actin was labeled with phalloidin (cyan). Scale bars, 5 μm and 1 μm. **(F, G)** Quantification of podosome size (μm²) (n≥158) and podosome number per cell (n≥21) were analyzed with ImageJ. Data are presented as mean ± SD from three independent experiments. Statistical significance was assessed using unpaired t test (C, D) or one-way ANOVA followed by Tukey multiple comparisons test (F, G). ns, not significant; **, P<0.01; ***, P<0.001. **(H)** Schematic model of podosome maturation promoted by Tks5 and Dyn2-Y597 phosphorylation.

Given our earlier observation that *C. albicans* infection promotes the association of Dyn2 with podosomes, we asked whether elevating Tks5 levels to drive maturation directly alters the spatial distribution of Dyn2. Co-transfection of Tks5-mEGFP and Dyn2-mCherry in RAW 264.7 macrophages revealed that Tks5 overexpression markedly enhanced Dyn2 recruitment to the podosome core, where Tks5 colocalized precisely with F-actin (Fig. EV7). Crucially, this enhanced recruitment was attenuated upon transfection of a Tks5 mutant lacking the two SH3 domains responsible for Dyn2 binding (Tks5ΔSH3AE). This cellular dependency beautifully correlates with our *in vitro* biochemical data, demonstrating that interaction with Tks5 stabilizes Dyn2 within podosomes.

We next investigated whether Dyn2-Y597 phosphorylation also actively drives podosome maturation. To address this, we expressed phospho-mimetic Dyn2Y597E or phospho-deficient Dyn2Y597F mutants in RAW 264.7 macrophages. Strikingly, expression of Dyn2Y597E increased podosome size and promoted the enrichment of Dyn2 into prominent ring-like structures that encircle the podosome core, whereas Dyn2Y597F reduced both podosome size and number (Fig. 6E-G). Collectively, these results provide *in vivo* validation of our biochemical model, demonstrating that Dyn2-Y597 phosphorylation and Tks5 interaction act as a molecular switch, transitioning Dyn2 into a structural actin bundler that dictates macrophage podosome maturation (Fig. 6H).

### Podosome maturation enhances macrophage mechanical immunity against *C. albicans*

Having demonstrated that *C. albicans* infection drives podosome maturation with Dyn2 phosphorylation and Tks5 as its structural engine, we finally asked whether podosome maturation is required for macrophage mechanical immunity against *C. albicans*. Using our Tks5-OE macrophages to artificially drive podosome maturation, we evaluated its effect on phagocytic efficiency. Macrophages were incubated with opsonized GFP-expressing *C. albicans* for 30 minutes to quantify engulfment (Fig. 7A). Tks5-OE cells exhibited a ∼2-fold increase in internalized fungal volume compared to controls (Fig. 7B, C), indicating that induced podosome maturation is sufficient to enhance phagocytic capacity.

**Fig. 7.**
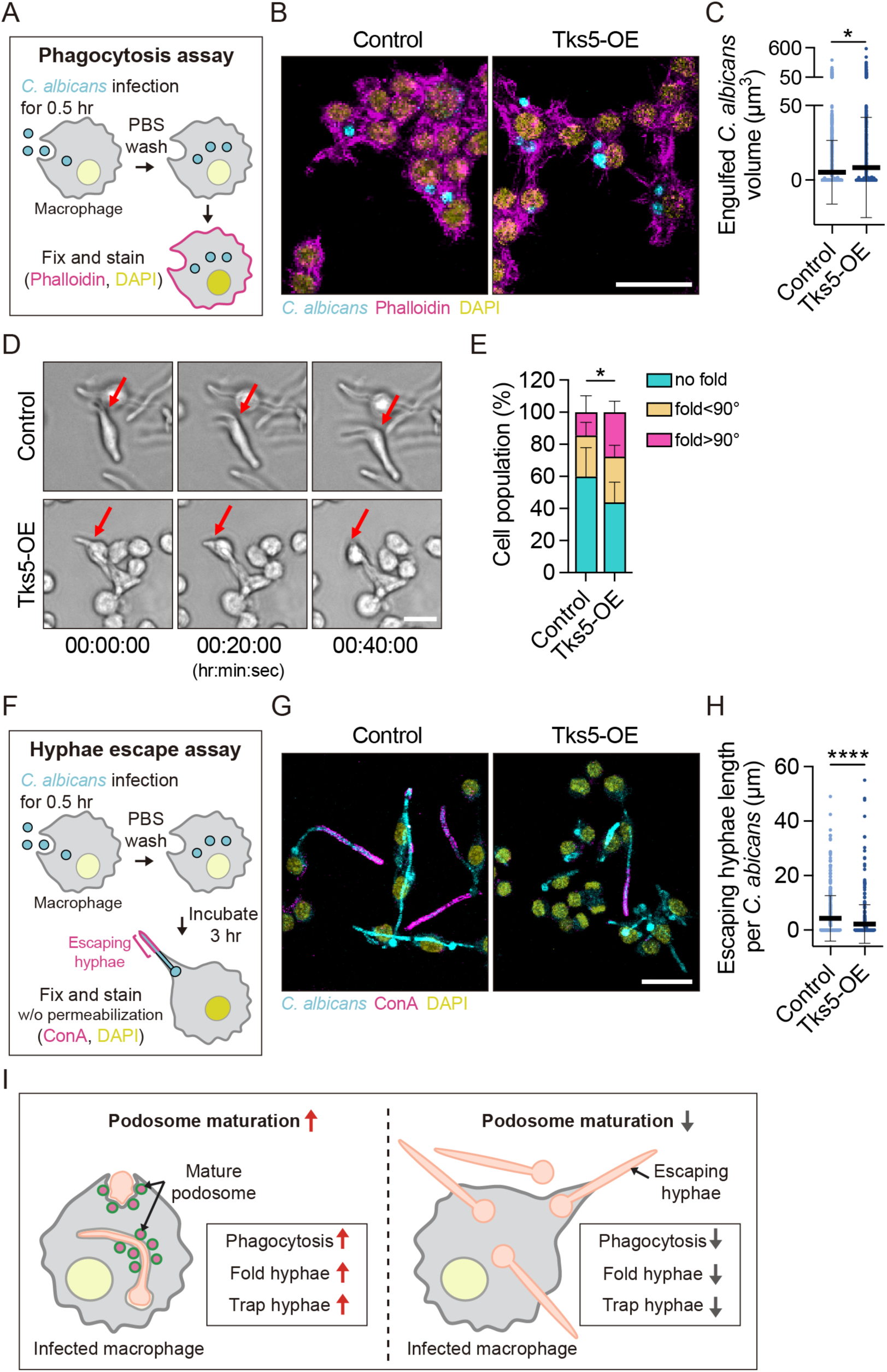
Tks5-mediated podosome maturation promotes macrophage mechanical immunity against *C. albicans*. **(A)** Workflow of the phagocytosis assay to evaluate phagocytic efficiency. **(B, C)** Representative fluorescence images (B) and volumetric quantification (C, n≥1199) of internalized GFP-*C. albicans* (cyan) in control and Tks5-OE RAW 264.7 cells. F-actin and nuclei were visualized with phalloidin (magenta) and DAPI (yellow), respectively. Scale bar, 20 µm. **(D, E)** Time-lapse microscopy (D) and quantification (E, n=3) of hyphae folding dynamics. Red arrows indicate active folding sites. Statistical significance was determined using contingency analysis followed by the Chi-square test. *, P<0.05. Scale bar, 20 µm. **(F)** Workflow of the hyphae escape assay. **(G, H)** Fluorescence images (G) and quantification of escaping hyphae length (H, n≥357). Extracellular hyphae were labeled with ConA (magenta). Scale bar, 20 µm. Data are presented as mean ± SD from three independent experiments. Statistical significance was assessed using two-way ANOVA followed by Tukey multiple comparisons test. *, P<0.05; ****, P<0.0001. **(I)** Model of podosome maturation in macrophage antifungal defense.

The hyphae form of *C. albicans* is a key virulence determinant, enabling tissue invasion and facilitates escape from macrophages^46–48^. To counteract this, macrophages utilize specialized mechanical strategies to physically fold hyphae and restrict outgrowth^26^. To determine whether podosome maturation influences this folding ability, we performed live-cell imaging of macrophages incubated with live *C. albicans* for 6 hours. Folding behavior was categorized into three groups: no fold, moderate fold (angles < 90°), and strong fold (angles > 90°). Tks5-OE macrophages demonstrated superior mechanical capacity, exhibiting a lower percentage of cells with no folding activity and a significantly higher frequency of strong folding events (Fig. 7D, E). Contingency analysis revealed significant shifts in folding capacity between Tks5-OE and control cells.

To determine if this enhanced physical deformation effectively limits fungal escape, we performed a hyphae escape assay (Fig. 7F). Following 30 min of internalization, washing, and a 3-hour incubation, GFP expressing fungi with extracellular escaping hyphae were stained with concanavalin A (ConA), which binds mannose-rich glycoproteins exposed exclusively on hyphae that have breached the cell membrane^49,50^. Tks5-OE macrophages significantly restricted hyphae growth, resulting in reduced escaping hyphae length (Fig. 7G, H). Together, these gain-of-function findings demonstrate that driven podosome maturation is sufficient to efficiently engulf, fold, and trap *C. albicans*.

To definitively establish whether podosome maturation is a strict physiological requirement for these processes, we generated Tks5-knockout (KO) RAW 264.7 macrophages using CRISPR/Cas9 (Fig. EV8A). While Tks5-KO macrophages maintained baseline podosome morphology (Fig. EV8B-D), they exhibited severe defects across all mechanical defense metrics: fungal engulfment was reduced (Fig. EV8E, F), hyphae folding was impaired (Fig. EV8G, H), and escaping hyphae were significantly longer (Fig. EV8I, J). Furthermore, immunofluorescence revealed that endogenous Tks5 localized specifically to phagocytic cups and phagosome-associated F-actin networks during *C. albicans* infection (Fig. EV8K). Altogether, these findings support a model in which podosome maturation provides the mechanical resilience required for robust macrophage immunity against *C. albicans* (Fig. 7I).

## Discussion

In this study, we show that *C. albicans* infection drives the maturation of macrophage podosomes into mechanically resilient structures that enhance antifungal defense (Fig. 8). Resting macrophages predominantly harbor nascent podosomes, whereas pathogen exposure induces a coordinated maturation program marked by upregulation of Tks5, Src, MMP14, and Cortactin. This response is consistent with previous reports of infection-induced Tks5 transcription^51^, LPS-dependent Src activation^52^, and phosphotyrosine enrichment at phagocytic podosomes^29^. Together with our gain- and loss-of-function analyses, these findings demonstrate that podosome maturation provides the mechanical strength required to engulf, constrain, and trap fungal hyphae. More broadly, similar induction of Tks5 during osteoclastogenesis^53^ and pathogenic T-cell activation^54^ suggests that regulated podosome maturation is a conserved strategy for scaling cellular mechanical output in response to physiological demand.

**Fig. 8.**
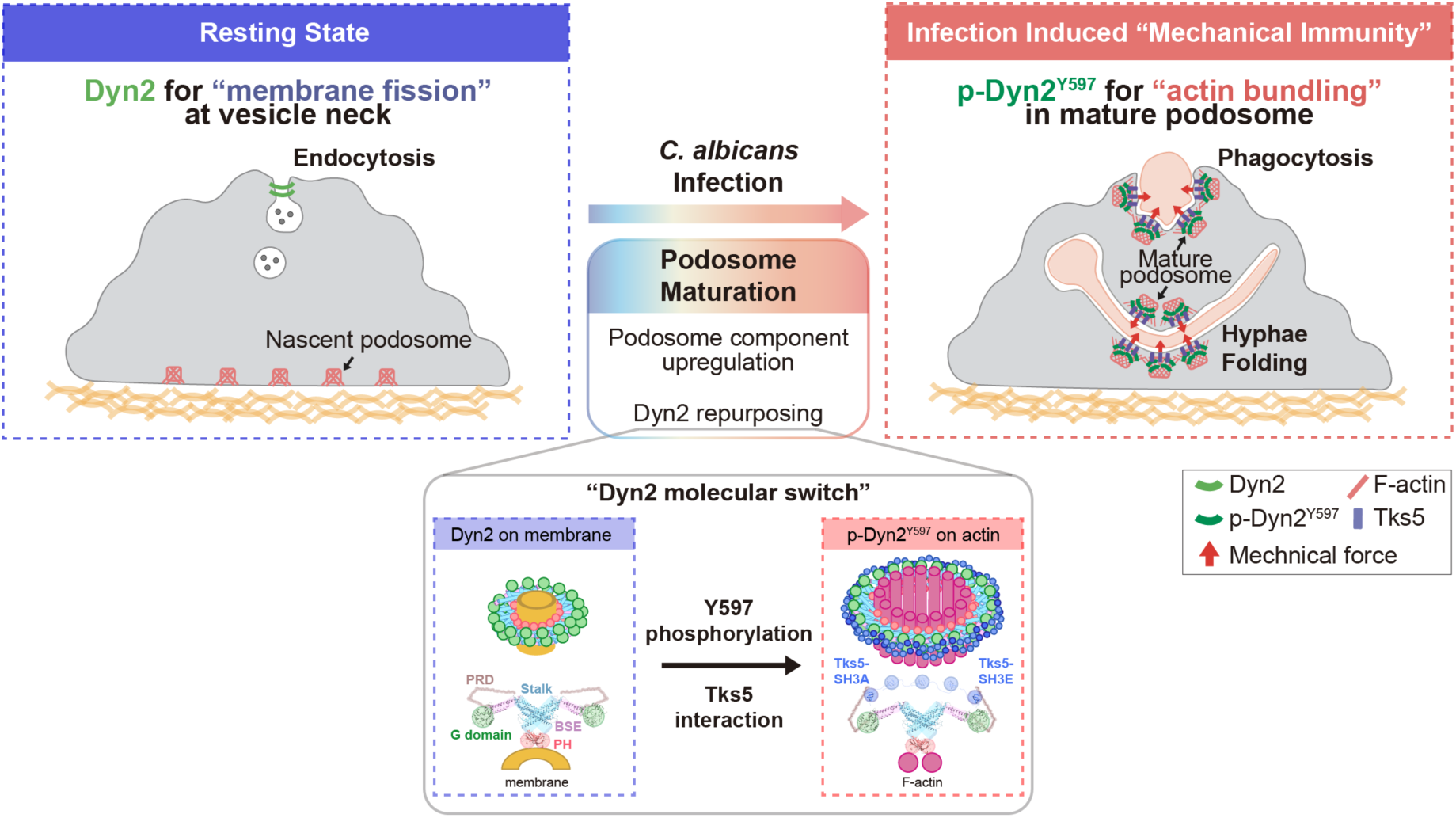
Model of *C. albicans* infection-induced podosome maturation for macrophage mechanical immunity. In their resting state, macrophages primarily harbor nascent podosomes and utilize Dyn2 for membrane fission during endocytosis. Upon exposure to pathogenic stimuli, such as *C. albicans*, the upregulation of key podosome components, including Tks5 and Src, drives podosome maturation. During this transition, Dyn2 undergoes a functional switch: Src-mediated phosphorylation at Y597 triggers its translocation from the plasma membrane to the F-actin cytoskeleton. Tks5 then interacts with this phosphorylated Dyn2 through its SH3 domains, promoting the assembly of Dyn2 helices that encircle and bundle actin filaments to facilitate podosome maturation. Collectively, this maturation process is critical for the efficient phagocytosis of *C. albicans*, enabling macrophages to physically fold and constrain fungal hyphae, thereby preventing pathogen escape and ensuring containment.

At the molecular level, the enhanced mechanical resilience of mature podosomes depends on the actin-bundling activity of Dyn2^33,35^. Here, we identify a cooperative molecular switch that repurposes Dyn2 from a membrane-remodeler into an actin bundler (Fig. 8, bottom). Src-mediated phosphorylation of Dyn2 at Y597 biases its activity toward the actin cytoskeleton, while Tks5 promotes Dyn2 assembly and stabilizes its association with actin through a dynamic “catch-and-release” cycle. This mechanism resolves how a GTPase optimized for rapid membrane remodeling can be incorporated into long-lived yet dynamic podosomes. Structurally, Dyn2 assembles into large helices that encircle multiple actin filaments, resembling but distinct from its classical membrane-bound helices^55–57^. This architecture differs from the previously proposed model of internal Dyn2 helices formed under low-salt conditions^35^, likely due to the salt sensitivity of Dyn2 assembly, and underscores the need for future high-resolution structural studies using in situ cryo-electron tomography under strictly physiological conditions.

The capacity of immune cells to sense and adapt to their physical environment is critical for effective immunity^58^. Macrophage function is tightly governed by microenvironmental mechanics, where matrix stiffness, shear stress, and cyclic stretch heavily influence polarization, migration, and phagocytosis^59–64^. As highly mechanosensitive structures, podosomes are central to these responses during macrophage migration and pathogen engulfment^11,24–29^. By demonstrating that macrophage rapidly restructure and repurpose podosomes to physically overpower invading fungi, this work establishes a novel paradigm of mechanical immunity. These insights provide a molecular foundation for understanding the biomechanical regulation of host defense and inflammatory disease.

## Methods

### Isolation and Culture of Primary Mouse Bone Marrow-Derived Macrophages (mBMDMs)

BMDMs were isolated from C57BL/6 mice euthanized using CO_2_. The hind limbs were sterilized with 75% ethanol, and the skin and muscles near the joints of the femur and tibia were carefully removed. The femur and tibia were separated at the joints, and surrounding muscle tissue was cleared to expose the bones. Both bones were then placed in PBS containing 2% heat-inactivated FBS. Bone marrow was flushed out using a 5 ml syringe filled with 2% FBS in PBS and collected in sterile tubes. The marrow suspension was homogenized by gentle pipetting and centrifuged at 500 xg for 4 minutes. Red blood cells were lysed using RBC lysis buffer, followed by the addition of 2% FBS in PBS. After another round of centrifugation, the cells were washed three times with 2% FBS in PBS. To induce differentiation, the cells were plated in 10 cm dishes and cultured for seven days in complete DMEM, containing high-glucose DMEM (HyClone), 10% heat-inactivated FBS (Corning), and 1 mM sodium pyruvate (Gibco), supplemented with 30% L929 cell-conditioned medium. Finally, the cells were incubated in complete DMEM supplemented with 10 ng/ml M-CSF for 24 hours.

### RAW 264.7 Cell Culture

RAW 264.7 cell, a murine macrophage-like cell line derived from an Abelson murine leukemia virus-induced tumor, were generously provided by Dr. Li-Chung Hsu’s laboratory. The cells were cultured in high-glucose DMEM supplemented with 10% heat-inactivated FBS, 1 mM sodium pyruvate, and 1x antibiotic-antimycotic solution (Gibco). For phagocytosis experiments, the culture medium was replaced with phagocytosis medium containing high-glucose DMEM, 10% FBS, and 1 mM sodium pyruvate.

### Candida albicans (C. albicans) Culture

The *C. albicans* strain SC5314 and OG-1, which expresses GFP, was obtained from Dr. Li-Chung Hsu’s laboratory. The strain was cultured on YPD agar slant tubes for overnight at 30°C prior to use.

### Observation of Podosome Morphology

For observing podosome morphology in mBMDMs, mBMDMs were cultured in complete DMEM supplemented with 10 ng/mL M-CSF and seeded overnight onto fibronectin-coated coverslips. Cells were either fixed immediately to serve as the uninfected control (0 h) or challenged with *C. albicans*. For infection timelines, macrophages were incubated with the fungus for 0.5 h and then either fixed immediately (0.5 h sample) or washed with PBS to remove unengulfed fungi and further incubated for an additional 1.5 h (2 h sample) or 3.5 h (4 h sample) prior to fixation. Following the respective time courses, cells underwent immunofluorescence staining. Images were acquired using an LSM700 confocal microscope (Carl Zeiss). For RAW 264.7 macrophages, cells were cultured overnight at 37 °C in a 5% CO₂ incubator. The following day, cells were reseeded onto fibronectin-coated coverslips and incubated for 3 hours under the same conditions. Podosome morphology was then examined by immunofluorescence staining, and images were acquired using LSM700 or LSM980 confocal microscope (Carl Zeiss).

### Immunofluorescence Staining

Cells were washed with PBS and fixed with 4% paraformaldehyde (#P6148, Sigma-Aldrich) in PBS for 10 minutes at room temperature. After fixation, cells were permeabilized using 0.1% Triton X-100 in PBS for 15 minutes. Blocking was performed with a solution containing 3% bovine serum albumin (BSA), 5% normal donkey serum (NDS), and 0.01% saponin in PBS for 1 hour at room temperature. Cells were subsequently incubated with the indicated primary antibodies, followed by fluorophore-conjugated secondary antibodies for detection. Antibodies used in this study are listed in Table S1.

### Western Blot

Cells were seeded in 6 cm dishes and harvested the following day. After washing with PBS, cells were lysed using RIPA buffer with 1 mM Na₃VO₄, 50 mM NaF, 1x protease inhibitor (#04693116001, Roche), and 1x phosphatase inhibitor (#04906837001, Roche). Lysates were incubated on ice and centrifuged at 15,000 rpm for 10 minutes at 4°C. The supernatant was collected, and protein concentrations were measured using the Bio-Rad protein assay. Equal amounts of protein were mixed with SDS loading buffer, heated at 95°C for 10 minutes, and separated on SDS-PAGE gels at 120 V for 90 minutes. Proteins were transferred to PVDF membranes at 100 V for 90 minutes at 4°C. Membranes were blocked with 3% BSA in TBST for 1 hour at room temperature and incubated with primary antibodies overnight at 4°C. After washing, membranes were incubated with HRP-conjugated secondary antibodies for 30 minutes at room temperature. Protein bands were detected using chemiluminescence. A full list of antibodies is provided in Table S1.

### Protein Purification

To express and purify dynamin, human dynamin was subcloned into the pIEX6 expression vector (Novagen) and subsequently expressed in Sf9 insect cells. Dynamin was purified using a GST-fused SH3 domain from amphiphysin-2, followed by overnight dialysis. The final protein was stored in a buffer composed of 1 mM DTT, 150 mM KCl, 1 mM EGTA, 20 mM HEPES (pH 7.5), and 10% glycerol^65^.

For Tks5 expression, full-length human Tks5 (SH3PXD2A) and various truncated constructs were cloned into either pGEX4T-1 or pET-15b for expression in *E. coli*. Tks5 proteins were purified using glutathione Sepharose beads (GE Healthcare) or Ni-NTA resin (Qiagen), followed by elution according to the manufacturer’s instructions. Tks5 was subjected to overnight dialysis and kept in buffer containing 20 mM HEPES, 150 mM KCl, 1 mM DTT, and 10% glycerol. Plasmids used in this study is provided in Table S2.

### F-actin Bundle Sedimentation Assay

The actin bundling assay followed protocols adapted from previous reports^33,34^. In brief, 10 µM rabbit skeletal muscle actin (#AKL99-C, Cytoskeleton) was diluted in a general actin buffer (5 mM Tris, pH 7.4, 0.2 mM ATP, 0.2 mM CaCl₂, 0.5 mM DTT) and incubated at 4°C for 1 hour. Aggregates were removed by centrifugation at 20,000 xg for 15 minutes. The resulting G-actin supernatant was then polymerized into F-actin by adding polymerization buffer (2.5 mM Tris, pH 7.4, 0.1 mM CaCl₂, 0.25 mM DTT, 50 mM KCl, 2 mM MgCl₂, 1 mM ATP) and incubating at room temperature for 1 hour. To assess Dyn2-mediated bundling, 5 µM F-actin was incubated with 1 µM Dyn2, with KCl adjusted to 75-150 mM. After 30 minutes at room temperature, samples were centrifuged at 14,000 xg for 20 minutes. Proteins in the supernatant and pellet were solubilized in SDS sample buffer, separated by SDS-PAGE, stained with Coomassie blue, and quantified using ImageJ. Bundling activity was calculated as the ratio of actin in the pellet to total actin.

### GST Pull-Down Assay

Glutathione beads carrying 40 μg GST or GST-Tks5 were incubated with 2 μg purified His-Dyn protein in 1 ml binding buffer (PBS containing 1% Tween-20, 1 mM DTT, and 10% glycerol) at 4 °C for 1 hour. After incubation, the beads were washed three times with binding buffer supplemented with 1.2% Tween-20. The proteins bound on the beads were then analyzed by Western blot.

### Transmission Electron Microscopy (TEM)

To observe Dyn2-actin bundles, 5 μM F-actin was incubated with or without 1 μM Dyn2 and 1 μM mini-Tks5 for 30 minutes at room temperature. Following incubation, the samples were applied to carbon-coated, glow-discharged EM grids and stained with 2% uranyl acetate. Imaging was performed using a Hitachi H-7650 transmission electron microscope operating at 75 kV.

### GTPase Activity Assay

Purified Dyn2 (1 μM) and actin (5 μM) bundles were prepared as described above by incubating the proteins at room temperature for 30 minutes, with or without mini-Tks5. The reactions were then warmed to 37°C, and GTP was added to a final concentration of 1 mM with gentle pipetting. GTP hydrolysis was monitored over time using a colorimetric malachite green assay to detect the release of inorganic phosphate.

### Liposome Binding, TEM, and GTPase Activity Assays

The lipid composition used in this study consisted of DOPC, DOPS, and PI(4,5)P₂ in a molar ratio of 70:15:15. To generate liposomes, lipid mixtures were first dried and then rehydrated in a buffer containing 150 mM KCl and 20 mM HEPES (pH 7.5). The suspension underwent three freeze-thaw cycles before being extruded through a 0.1-μm polycarbonate membrane (Whatman) using an Avanti mini-extruder.

To assess membrane-binding capacity, 0.5 μM Dyn2 and/or 0.5 μM mini-Tks5 were incubated with 150 μM of 100-nm liposomes for 30 minutes at room temperature. Following incubation, liposome-bound proteins were pelleted by centrifugation at 15,000 xg and resuspended in assay buffer (20 mM HEPES, 150 mM KCl). Supernatants containing unbound proteins were collected separately. Both pellet and supernatant fractions were analyzed by SDS-PAGE followed by Coomassie staining. Protein band intensities were quantified using ImageJ, and membrane-binding efficiency was calculated as the ratio of pellet-associated protein to the total protein input.

For electron microscopy, 1 μM Dyn2 and/or 1 μM mini-Tks5 were mixed with 25 μM liposomes and incubated for 30 minutes at room temperature. The mixture was applied to glow-discharged, carbon-coated grids for 5 minutes, stained with 2% uranyl acetate for 2 minutes, and imaged using a Hitachi H-7650 transmission electron microscope operated at 75 kV.

To measure GTPase activity, 0.5 μM purified Dyn2 was incubated with or without 0.5 μM mini-Tks5 in buffer containing 150 μM liposomes, 150 mM KCl, 20 mM HEPES, 1 mM MgCl₂, and 1 mM GTP at 37 °C. GTP hydrolysis was monitored using a colorimetric malachite green assay to quantify released inorganic phosphate.

### Fluorescence Visualization of Dyn2-Actin Bundling Activity

Dyn2-actin bundles were assembled by incubating 5 µM F-actin with 1 µM 25% of Dyn2 labeled with Alexa Fluor 488 (Dyn2-488) and/or 1 µM mini-Tks5 as described above. To monitor the Dyn2 dynamic following GTP addition, pre-formed Dyn2-488-actin bundles were spiked with a solution containing GTP and Alexa Fluor 594-labeled Dyn2 (Dyn2-594). At specific time points (1 and 5 minutes), the reaction mixtures were diluted tenfold in an imaging buffer (10 mM HEPES pH 7.4, 75 mM KCl, 1 mM MgCl_2_, 0.1 mM EDTA, 1 mM DTT, 3 mg/ml glucose, 2 mg/ml BSA, and 0.2% methylcellulose, 50 μl/ml phalloidin). The samples were immediately dropped on slide and covered with coverslip. The images were taken by LSM700 confocal microscope (Carl Zeiss).

### Cryo-EM Imaging and Data Processing

Actin bundles were assembled by incubating 0.5 µM F-actin with 0.5 µM Dyn2, 0.5 µM mini-Tks5, and 1 mM GMPPCP for 30 min at room temperature. Following assembly, 0.05% Tween-20 was added for 5 min. Bundles were sedimented by centrifugation at 2,000 xg for 1 min. 8 µl of the sedimented fraction was applied to glow-discharged holey carbon grid (Quantifoil Cu R2/2, 200 mesh). Grid was blotted for 2 s at 22 °C and 90% humidity using a Leica EM GP2 Automatic Plunge Freezer and vitrified in liquid ethane. Data were acquired on a Glacios Cryo-TEM (Thermo Fisher Scientific) operating at 200 kV equipped with a Falcon 4 direct electron detector. Movies were collected in counting mode using EPU software at a magnification of 73,000x, resulting in a pixel size of 1.9 Å. Each movie comprised 40 frames with a total dose of 30 electrons per Å^2^. All data processing was performed in CryoSPARC. Micrographs underwent patch motion correction and patch CTF estimation. Dyn2-actin bundles were identified using the CryoSPARC manual picker, followed by particle extraction and multiple rounds of 2D classification to isolate classes clearly exhibiting Dyn2 helical structures encircling actin filaments.

### Dyn2-Actin Bundle Straightness Assay

Dyn2-actin bundles were assembled by incubating 5 µM F-actin with 1 µM Dyn2 and/or 1 µM mini-Tks5 as previously described. For imaging, 1 µl of the Dyn2-actin mixture was diluted with 9 µl of imaging buffer containing 10 mM HEPES (pH 7.4), 75 or 150 mM KCl, 50% Fluoromount-G (SouthernBiotech), and 1:500 phalloidin-Alexa Fluor 594. The diluted mixture was applied onto a coverslip and imaged using LSM700 confocal microscope (Carl Zeiss). Bundle straightness was quantified by calculating the ratio of the end-to-end distance to the total contour length of each Dyn2-actin bundle.

### Mechanical Disruption Assay

Dyn2-actin bundles were assembled by incubating 5 µM F-actin with 1 µM Dyn2 and/or 1 µM mini-Tks5 as described above. To assess mechanical stability, bundles were subjected to centrifugation at varying speeds. Following centrifugation, 10 µl of the Dyn2-actin mixture was mixed by pipetting with 10 µl of imaging solution containing 10 mM HEPES (pH 7.4), 75 mM KCl, 50% Fluoromount-G (SouthernBiotech), and 1:500 phalloidin-Alexa Fluor 594. The mixture was applied to a coverslip and imaged using LSM700 confocal microscope (Carl Zeiss). Mean F-actin fluorescence intensity of Dyn2-actin bundles was quantified using ImageJ.

### Generation of Tks5 Overexpressing (Tks5-OE) RAW 264.7 Cells by Lentivirus System

To generate Tks5-overexpressing RAW 264.7 cells, a lentiviral system was employed. HEK293T cells were seeded in 6 cm plates and cultured overnight to reach 70% confluency. The next day, transfection reagents, plasmids pCMV, pMD.G, pAS4.1w.Pbsd-aOn (a gift from Dr. Li-Chung Hsu) with or without Tks5, and Opti-MEM were mixed with TransIT X2 transfection reagent (#MIR 6010, Mirus Bio). After a 15-minute incubation at room temperature, the transfection mix was added to HEK293T cells. Following an 18-hour incubation at 37°C with 5% CO₂, the medium was replaced with virus collection medium (DMEM containing 10% FBS, 1% BSA). Viral supernatants were collected over 3 days, filtered (0.2 μm), and stored at -80°C. RAW 264.7 cells were infected with the thawed lentivirus in the presence of 8 μg/ml polybrene. After 24 hours, cells were selected with 10 μg/ml blasticidin for 2 days. Tks5 expression was induced by adding 10 μg/ml doxycycline.

### Generation of Tks5 Knockout (Tks5-KO) RAW 264.7 Cells by CRISPR/Cas9 System

To generate Tks5 knockout (KO) RAW 264.7 cells, the mouse Tks5 gene sequence was retrieved from NCBI, and pair of sgRNAs targeting the longest isoform were designed using CRISPick: sg-F: 5′-caccggagaagccacatccgcgacg-3′, sg-R: 5′-aaaccgtcgcggatgtggcttctcc-3′. The annealed oligos were cloned into the EDCPP lentiviral vector (a gift from Dr. Ping-Hung Chen). Lentiviruses carrying the sgRNA constructs were packaged and used to infect RAW 264.7 cells. Infected cells were selected with 4 μg/ml puromycin for 2 days, followed by Shield-1 treatment to induce Cas9 expression and gene editing. Single-cell clones were isolated via limiting dilution, and successful knockout was confirmed by western blot and DNA sequencing.

### Transfection

Cells at approximately 70% confluence were transfected with the indicated plasmids using TurboFect™ Transfection Reagent (Thermo Scientific), following the manufacturer’s protocol. The plasmids used in this study is provided in the Table S2.

### Phagocytosis Assay

Macrophages were seeded on coverslips and cultured overnight. *C. albicans* yeast cells were opsonized with rabbit anti-*C. albicans* IgG in PBS for 1 hour at room temperature with rocking. After opsonization, yeast cells were washed once and then incubated with macrophages in phagocytosis medium at a MOI of 3 in a 5% CO₂ incubator for 30 minutes. After infection, unengulfed pathogens were removed by washing cells three times with PBS. Cells were fixed with 4% paraformaldehyde for 10 minutes, permeabilized with 0.1% Triton X-100 for 15 minutes, and then stained with phalloidin and DAPI for 20 minutes. LSM700 confocal microscope (Carl Zeiss) was used to acquire the images.

### Hyphae Folding Assay

Macrophages were seeded on coverslips and cultured overnight. *C. albicans* were opsonized with rabbit anti-*C. albicans* IgG in PBS for 1 hour at room temperature, then washed with PBS. Opsonized yeasts were added to macrophages in pre-warmed phagocytosis medium at a MOI of 1. Live cell imaging was performed by AXIO Observer Z1 inverted fluorescence motorized microscope (Carl Zeiss) for 6 hours. Cells were categorized into three groups based on their ability to fold hyphae: no fold, folding at angles less than 90° (fold < 90°), and folding at angles greater than 90° (fold > 90°). The distribution of each group was quantified. The contingency analysis was used to compare control and Tks5-OE/KO cell’s folding ability.

### Hyphae Escape Assay

Macrophages were seeded on coverslips and cultured overnight. *C. albicans* yeast cells were opsonized with rabbit anti-*C. albicans* IgG in PBS for 1 hour at room temperature, then washed with PBS. Opsonized yeasts were added to macrophages in pre-warmed phagocytosis medium at a MOI of 3 and incubated for 30 minutes at 37 °C with 5% CO₂. After infection, cells were washed three times with PBS to remove unengulfed yeasts and incubated for an additional 3 hours to allow hyphae escape. Cells were then fixed with 4% paraformaldehyde for 10 minutes, and stained with concanavalin A (ConA) conjugated Alexa Fluor 594 (Invitrogen) to label extracellular hyphae and DAPI to stain nuclei. Images were acquired using LSM700 confocal microscope (Carl Zeiss). The length of escaping hyphae per individual hypha was measured based on the ConA signal.

### Fluorescence microscopy

For fixed samples, images were collected using an LSM700 or LSM980 confocal microscope with a 63x, 1.4-NA oil-immersion objective (Carl Zeiss). For time-lapse imaging, cells were imaged using AXIO Observer Z1 inverted fluorescence motorized microscope (Carl Zeiss).

### Image Analysis

Immunofluorescence images were analyzed using ImageJ (Fiji) software, Imaris, and ZEN (Carl Zeiss) software. A threshold was set for actin to quantify the area of podosomes. Phagocytosis images were analyzed in Imaris by generating 3D models of cells and pathogens using the Cell and Surface tools. Internalized pathogens were identified by thresholding negative distances to the cell surface, and pathogen volumes were quantified in Imaris. Escaping hyphae were measured using the spline tool in ZEN to determine hyphae length.

### Statistical Analysis

Quantitative data are presented as the mean ± SD from at least three independent experiments. Statistical analyses were performed using unpaired t test, one-way or two-way analysis of variance (ANOVA) followed by Tukey’s multiple comparisons test or Fisher’s LSD test, and contingency analysis followed by Chi-square test for trend. All analyses were conducted using GraphPad Prism. A p-value less than 0.05 was considered statistically significant and is indicated as follows: P<0.05 (*), P<0.01 (**), P<0.001 (***) and P<0.0001 (****).

## Acknowledgments

We thank the staff of the Technology Commons (College of Life Science), the Imaging Core Facility, and the Cryo-EM Core Facility (First Core Laboratory) at National Taiwan University for their technical support. This work was supported by National Science and Technology Council 113-2320-B-002-023-MY3 and National Taiwan University grant 114L7834.

## Author Contributions

Conceptualization: CW, YWL; Methodology: CW, TC, TYL, YWL; Investigation: CW, TC, TYL, YWL; Visualization: CW; Supervision: YWL; Writing—original draft: CW, YWL; Writing—review & editing: CW, YWL

## Competing Interest Declaration

Authors declare that they have no competing interests.

## Expanded View Figures

**Fig. EV1.**
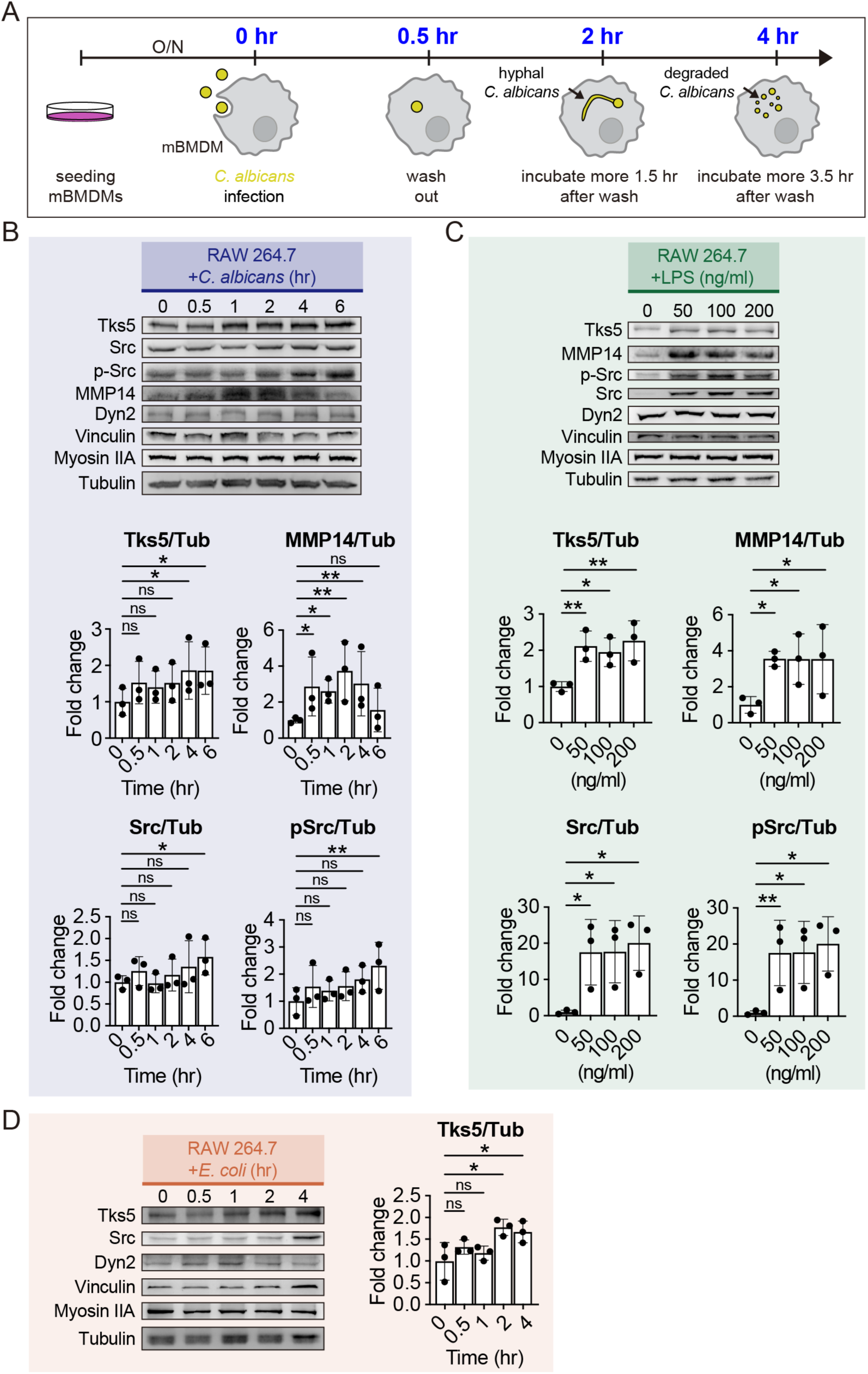
*C. albicans* infection promotes upregulation of podosome components in macrophages. **(A)** Schematic workflow of observing podosome morphology in mBMDMs after *C. albicans* infection. **(B)** Western blot and quantification of podosome components in RAW 264.7 cells infected with *C. albicans* for 0, 0.5, 1, 2, 4 and 6 hours at MOI=5. **(C)** Western blot and quantification of podosome components in RAW 264.7 cells added with 0, 50, 100 and 200 ng/ml LPS for overnight. **(D)** Western blot and quantification of podosome components in RAW 264.7 cells infected with *E. coli* for 0, 0.5, 1, 2, and 4 hours at MOI=10. Data are presented as mean ± SD of three independent experiments. Statistical significance was assessed using unpaired one-way ANOVA followed by the Fisher’s LSD test. ns, not significant; *, P < 0.05; **, P < 0.01.

**Fig. EV2.**
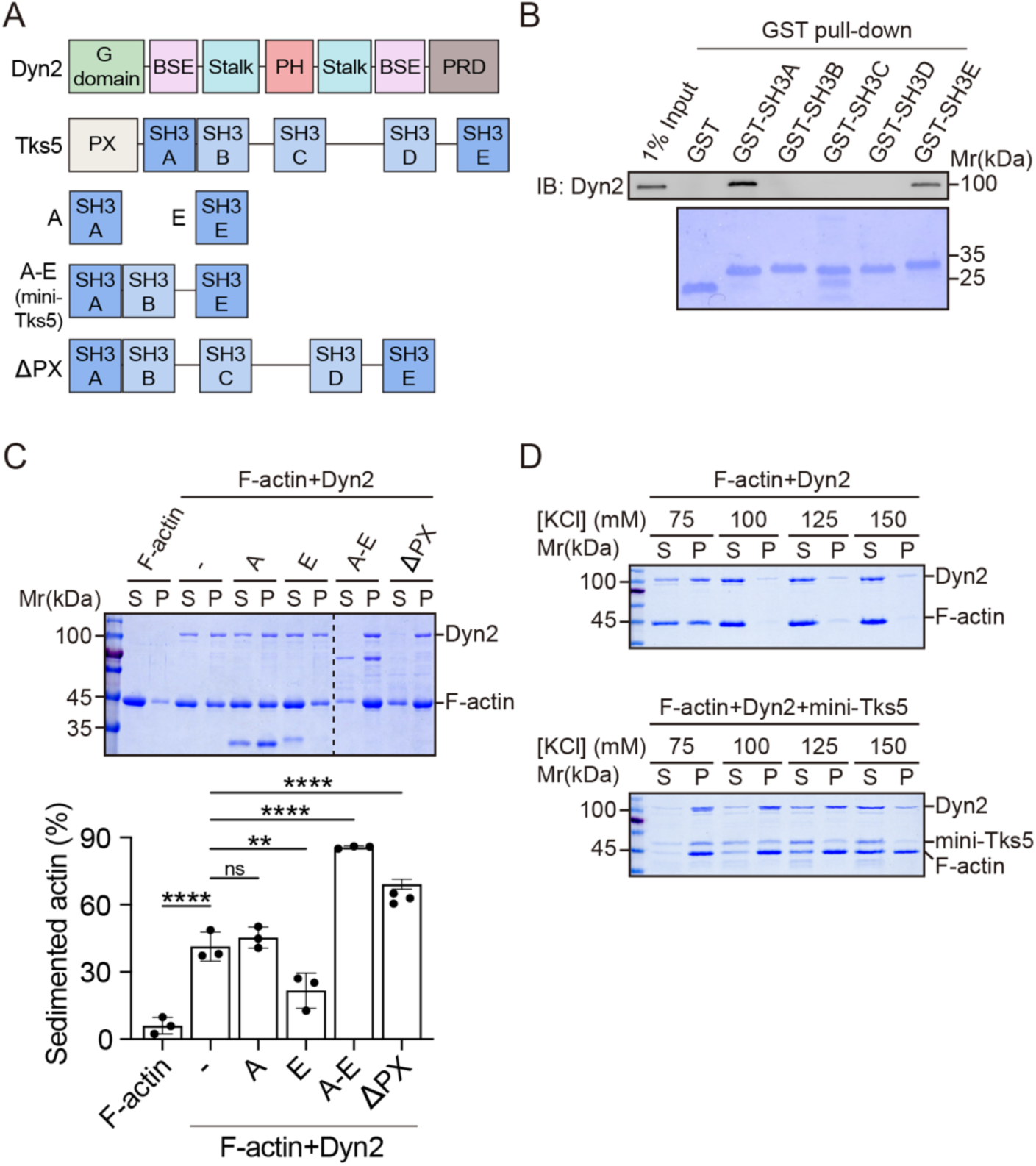
Tks5 interaction enhances Dyn2-mediated actin bundling. **(A)** Schematic diagram of Dyn2, Tks5, and various Tks5 SH3 constructs. **(B)** Direct binding between Dyn2 and individual SH3 domains of Tks5 was confirmed via GST pulldown assay. **(C)** Representative SDS-PAGE gel and quantification of F-actin bundle sedimentation assay in the presence or absence of various Tks5 SH3 constructs under 75 mM KCl. **(D)** Representative SDS-PAGE gels of F-actin bundle sedimentation assay in the presence or absence of mini-Tks5 under 75-150 mM KCl.

**Fig. EV3.**
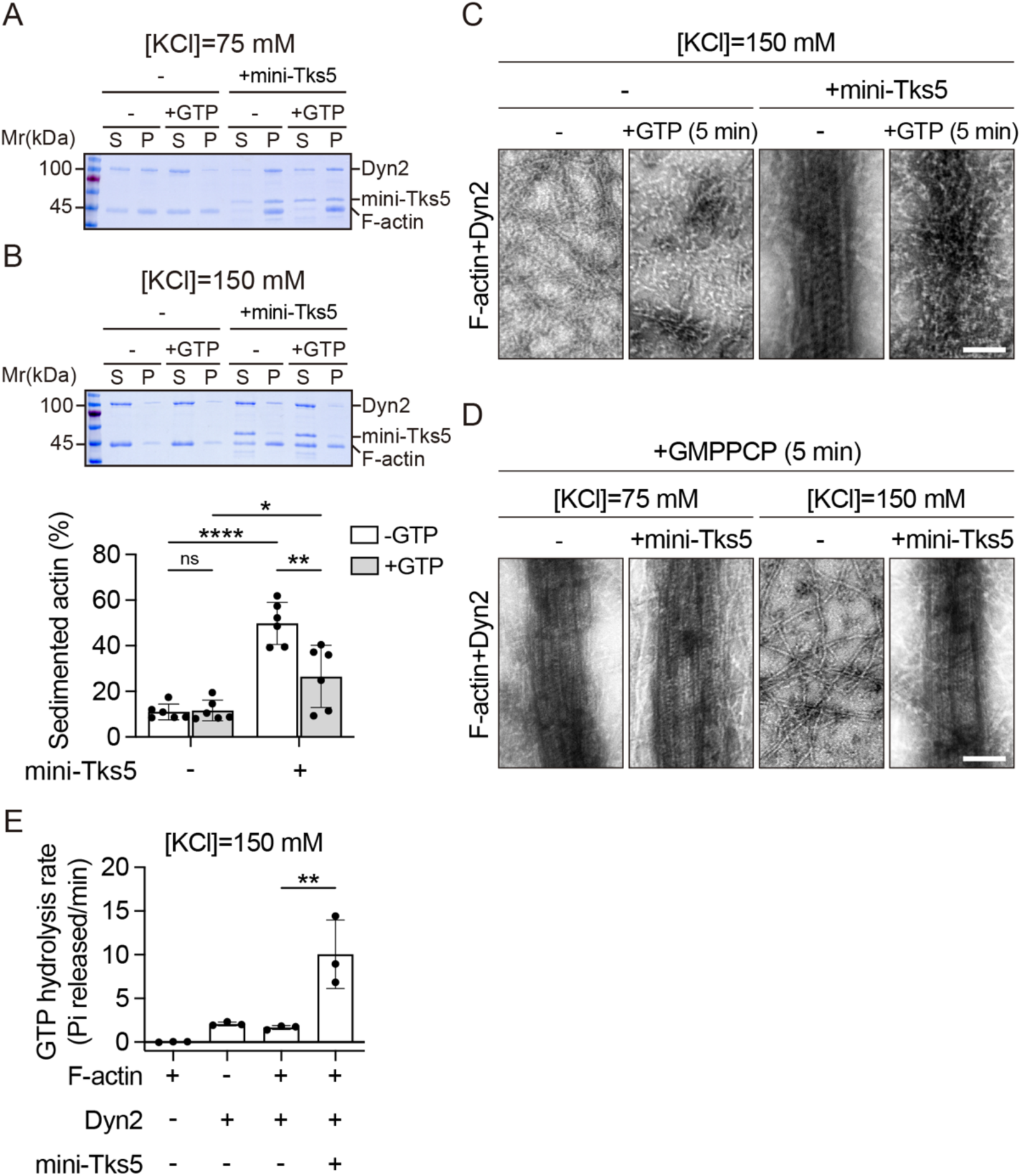
Tks5 reinforces Dyn2-actin bundles in the presence of GTP. **(A)** Representative SDS-PAGE gel of F-actin bundle sedimentation assay in the presence or absence of mini-Tks5 and GTP under 75 mM KCl. **(B, C)** F-actin sedimentation assays (B) and negative-stain TEM images (C) of Dyn2-actin bundles incubated with or without mini-Tks5 and GTP for 5 min at 150 mM KCl. **(D)** Negative-stain TEM images of Dyn2-actin bundles incubated with or without mini-Tks5 and GMPPCP for 5 min. Scale bars, 100 nm. **(E)** GTP hydrolysis rates of Dyn2-actin bundles in the presence or absence of mini-Tks5 at 150 mM KCl. Data are presented as mean ± SD from three independent experiments. Statistical significance was assessed using two-way (B) or one-way (E) ANOVA followed by Tukey multiple comparisons test. ns, not significant; *, P < 0.05; **, P < 0.01; ****, P < 0.0001.

**Fig. EV4.**
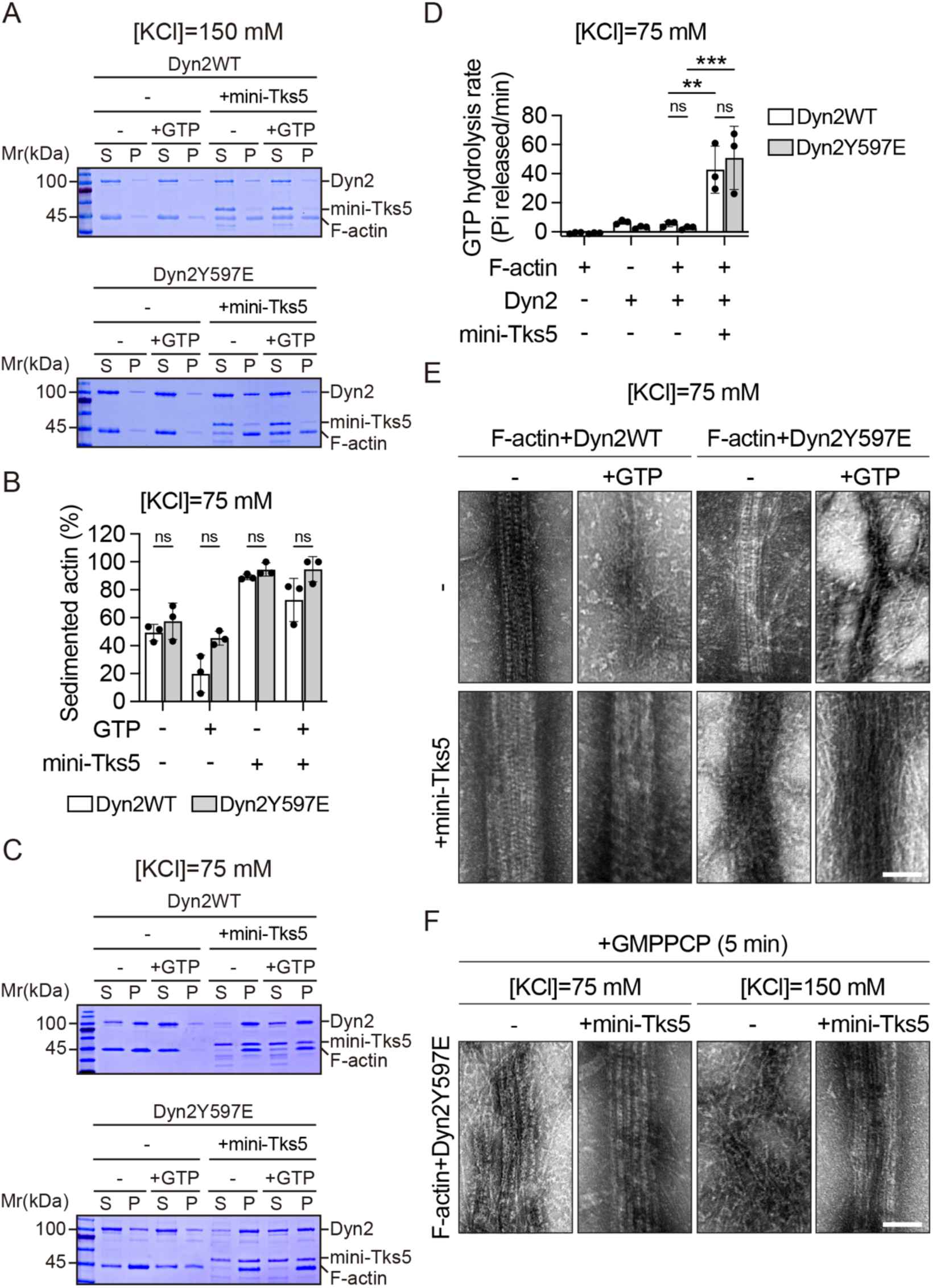
Dyn2 Y597 phosphorylation and Tks5 interaction cooperatively facilitate Dyn2-actin bundling. **(A)** Representative SDS-PAGE gels of F-actin bundle sedimentation assay using Dyn2WT or Dyn2Y597E in 150 mM KCl buffer, with or without mini-Tks5 and GTP. **(B, C)** Representative SDS-PAGE gels (B) and quantification (C) of F-actin bundle sedimentation assay using Dyn2WT or Dyn2Y597E in 75 mM KCl buffer, with or without mini-Tks5 and GTP. **(D)** GTP hydrolysis rate of Dyn2WT- or Dyn2Y597E-actin bundles with or without mini-Tks5 in 75 mM KCl buffer. Data are presented as mean ± SD from three independent experiments. Statistical significance was assessed using two-way ANOVA followed by Tukey multiple comparisons test. ns, not significant; **, P < 0.01; ***, P < 0.001. **(E)** Negative-stain TEM images of Dyn2WT- or Dyn2Y597E-mediated actin bundling with or without mini-Tks5 or GTP addition for 5 min in 75 mM KCl buffer. Scale bar, 100 nm. **(F)** Negative-stain TEM images of Dyn2Y597E-actin bundles incubated with or without mini-Tks5 and GMPPCP for 5 min. Scale bar, 100 nm.

**Fig. EV5.**
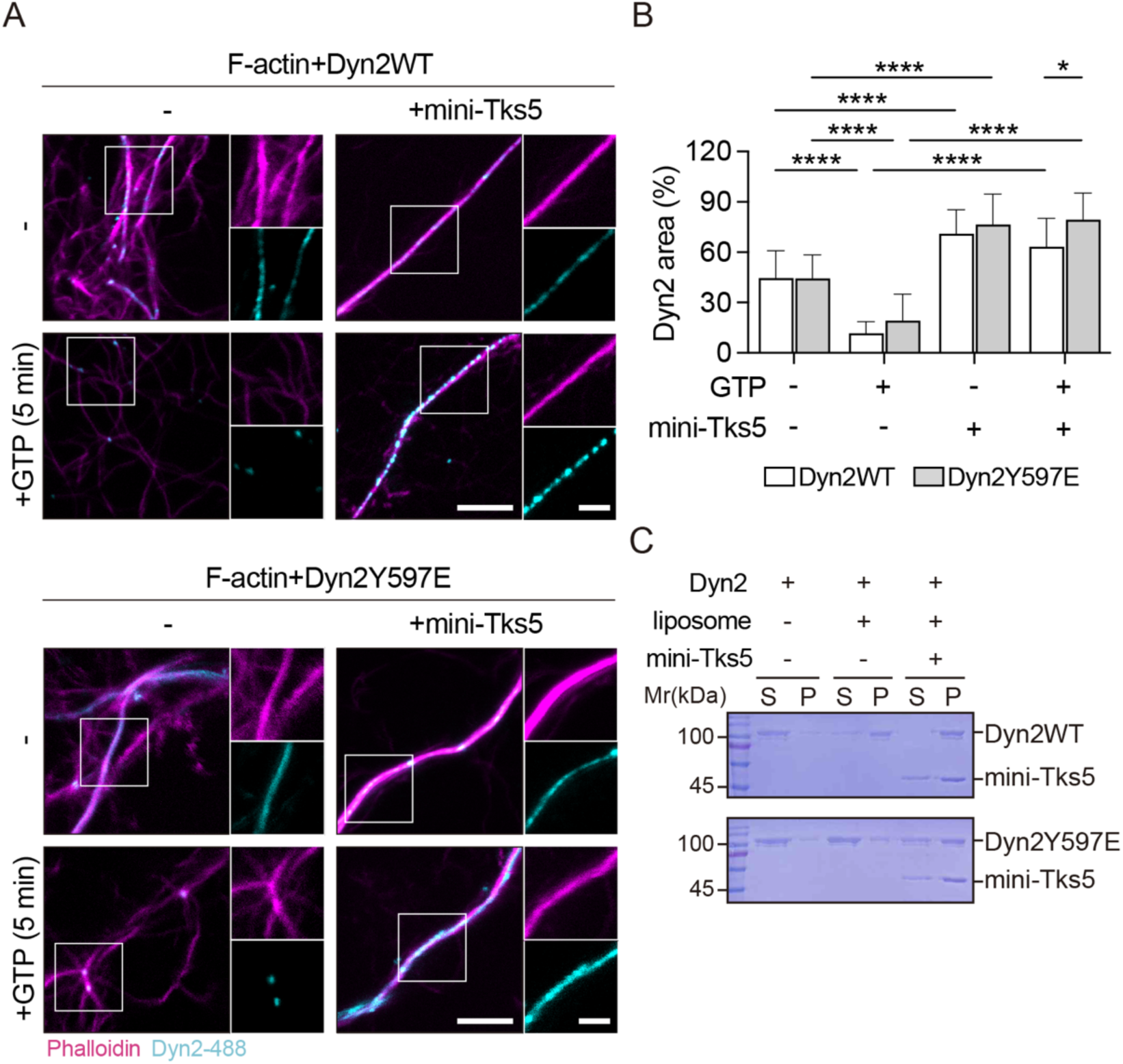
Dyn2 Y597 phosphorylation and Tks5 promotes Dyn2 assembly on actin. **(A)** Fluorescence imaging of Dyn2-actin bundling activity with or without mini-Tks5 and GTP in 75 mM KCl buffer. Scale bars, 5 µm and 2 µm. **(B)** Percentage of Dyn2 signal overlapping with actin, quantified using ImageJ (n≥14). Data are presented as mean ± SD from three independent experiments. Statistical significance was assessed using two-way ANOVA followed by Tukey multiple comparisons test. *, P < 0.05; ****, P < 0.0001. **(C)** Representative SDS-PAGE gels of liposome binding assay.

**Fig. EV6.**
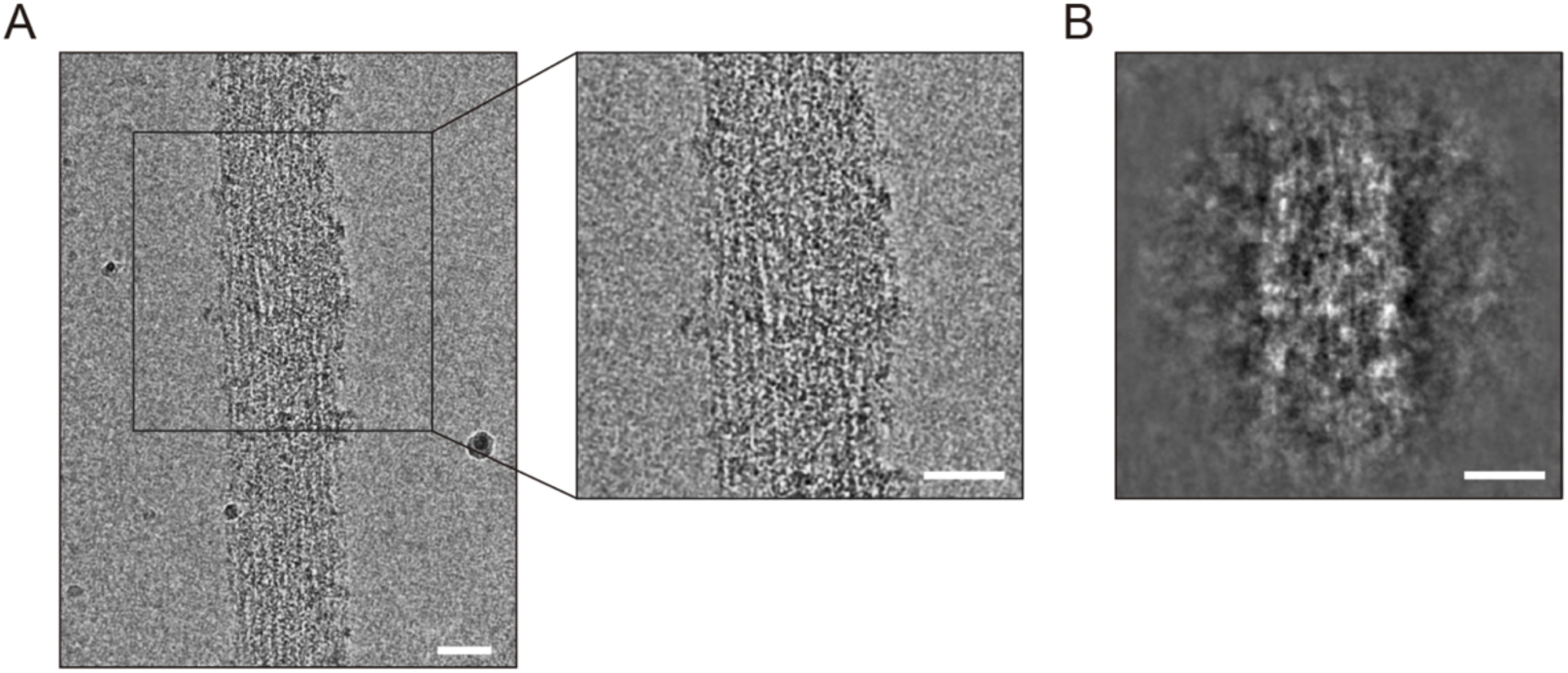
Cryo-EM analysis of Dyn2-actin bundles with mini-Tks5. **(A, B)** Representative cryo-EM image (A) and 2D class average (B) of Dyn2-actin bundle with mini-Tks5 and GMPPCP at 75 mM ionic strength condition. Scale bars, 50 nm.

**Fig. EV7.**
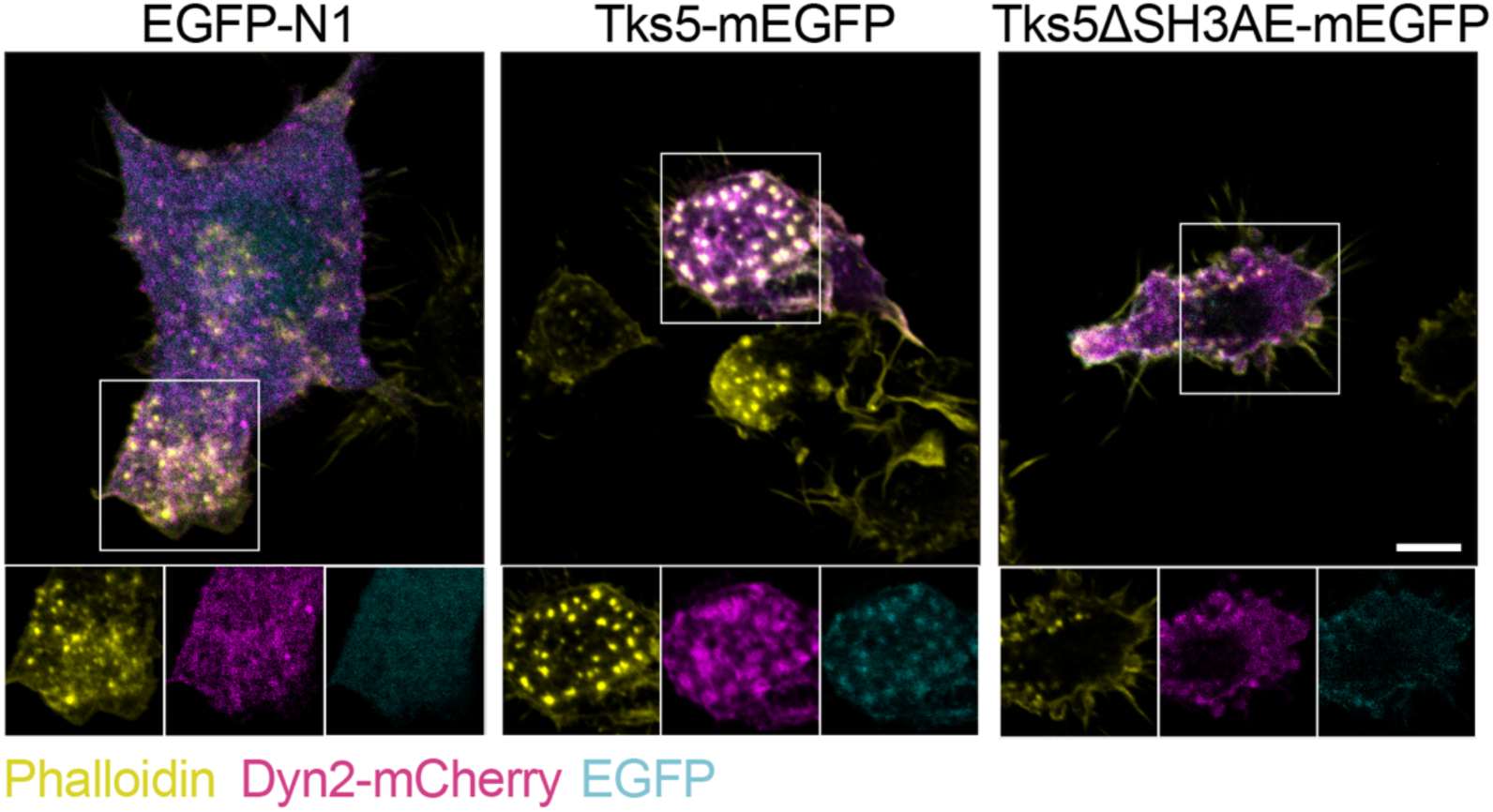
Tks5 overexpression promotes Dyn2 enrichment to macrophage podosomes. Fluorescence imaging of RAW 264.7 cells co-transfecting with Dyn2-mcherry (magenta) and EGFP-N1, Tks5-mEGFP or Tks5ΔSH3AE-mEGF (cyan). F-actin was labeled with phalloidin (yellow). Scale bar, 5 μm.

**Fig. EV8.**
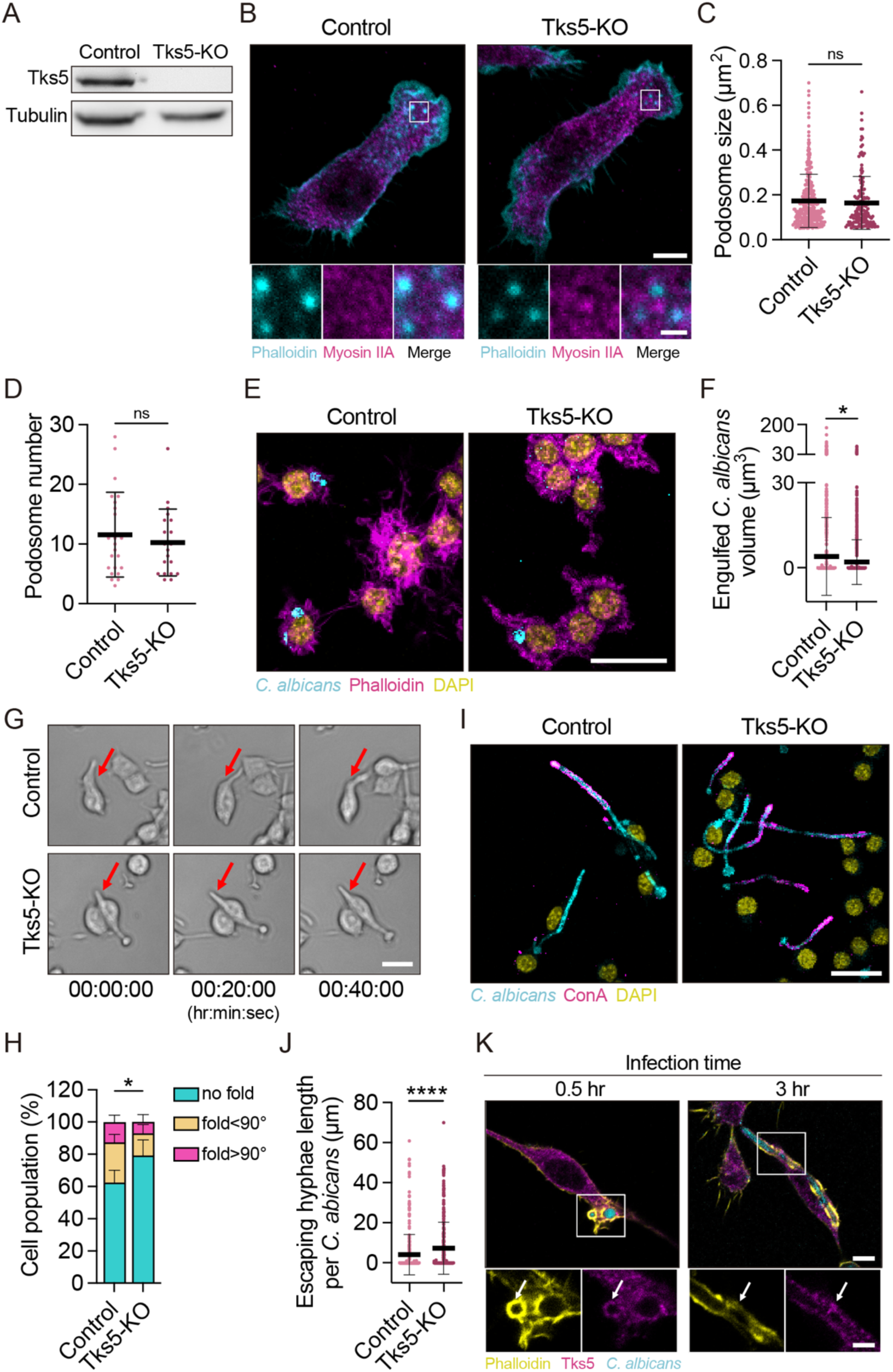
Tks5 depletion impairs macrophage mechanical immunity against *C. albicans*. **(A)** Western blot analysis confirmed the depletion of Tks5 in Tks5-KO cells compared to control cells. **(B)** Immunofluorescence staining of control and Tks5-KO RAW 264.7 cells. F-actin was labeled with phalloidin (cyan), and Myosin IIA was detected using specific antibodies (magenta). Scale bars, 5 μm and 1 μm. **(C, D)** Quantification of podosome size (μm²) (n≥161) and podosome number per cell (n≥20) were analyzed with ImageJ. **(E, F)** Representative fluorescence images (E) and volumetric quantification (F, n*≥*999) of internalized GFP-*C. albicans* (cyan) in control and Tks5-KO RAW 264.7 cells. F-actin and nuclei were visualized with phalloidin (magenta) and DAPI (yellow), respectively. Scale bar, 20 µm. **(G, H)** Time-lapse microscopy (G) and quantification (H, n=3) of hyphae folding dynamics. Red arrows indicate active folding sites. Statistical significance was determined using contingency analysis followed by the Chi-square test. *, P < 0.05. Scale bar, 20 µm. **(I, J)** Fluorescence images (I) and quantification of escaping hyphae length (J, n≥275). Extracellular hyphae were labeled with ConA (magenta). Scale bar, 20 µm. Data are presented as mean ± SD from three independent experiments. Statistical significance was assessed using one-way ANOVA followed by Tukey multiple comparisons test. ns, not significant; *, P < 0.05; ****, P < 0.0001. **(K)** Immunofluorescence staining of RAW 264.7 cells after 0.5 or 3 hours of *C. albicans* infection (MOI=3). Tks5 was stained by specific antibody (magenta), *C. albicans* expresses GFP (cyan), and F-actin was stained with phalloidin (yellow). Scale bars, 5 µm and 2.5 µm. White arrows indicate Tks5 localized around engulfed *C. albicans*.

**Table S1.** Antibodies used in this study.

| **Antibody** | **Source** | **Identifier** |
| --- | --- | --- |
| Tks5 | Santa Cruz | #30122 |
| MMP14 | Abcam | #ab51074 |
| Src | Cell Signaling | #2123S |
| p-Src | Cell Signaling | #6943S |
| Cortactin | Santa Cruz | #11408 |
| Dyn2 | Santa Cruz | #sc-6400 |
| Vinculin | Sigma-Aldrich | #V4505 |
| Paxillin | BD Biosciences | #610051 |
| Myosin IIA | Sigma-Aldrich | #M8064 |
| γ-tubulin | Sigma-Aldrich | #T6557 |
| *Candida albicans* Rabbit<br>Polyclonal Antibody | OriGene | #BP1006 |

**Table S2.** Plasmids used in this study.

| Plasmid | Vector | Source |
| --- | --- | --- |
| pIEx-6-Dyn2 | pIEx-6 | Schmid lab# |
| pIEx-6-Dyn2Y597E | pIEx-6 | (Lin et al., 2020) |
| pGEX-4T-Tks5-SH3A | pGEX-4T | In this study |
| pGEX-4T-Tks5-SH3B | pGEX-4T | In this study |
| pGEX-4T-Tks5-SH3C | pGEX-4T | In this study |
| pGEX-4T-Tks5-SH3D | pGEX-4T | In this study |
| pGEX-4T-Tks5-SH3E | pGEX-4T | In this study |
| pGEX-4T-Tks5-SH3A-E | pGEX-4T | In this study |
| pGEX-4T-Tks5-ΔPX | pGEX-4T | In this study |
| pET-15b-Tks5-SH3A-E | pET-15b | In this study |
| pAS4.1w-Pbsd-aOn-Tks5 | pAS4.1w-Pbsd-aOn | In this study |
| Tks5-mEGFP | mEGFP-N1 | In this study |
| Tks5ΔSH3AE-mEGFP | mEGFP-N1 | In this study |
| EDCPP-sgTks5 | EDCPP | In this study |
| Dyn2WT-mCherry | pmCherry-N1 | Addgene #27689 |
| Dyn2Y597E-mCherry | pmCherry-N1 | (Lin et al., 2020) |
| Dyn2Y597F-mCherry | pmCherry-N1 | (Lin et al., 2020) |

#Dr. Sandra Schmid in the Department of Cell Biology, University of Texas Southwestern Medical Center, Dallas, TX 75390, USA.

